# An atlas of gene regulatory networks for memory CD4^+^ T cells in youth and old age

**DOI:** 10.1101/2023.03.07.531590

**Authors:** Joseph A. Wayman, Alyssa Thomas, Anthony Bejjani, Alexander Katko, Maha Almanan, Alzbeta Godarova, Svetlana Korinfskaya, Tareian A. Cazares, Masashi Yukawa, Leah C. Kottyan, Artem Barski, Claire A. Chougnet, David A. Hildeman, Emily R. Miraldi

## Abstract

Aging profoundly affects immune-system function, promoting susceptibility to pathogens, cancers and chronic inflammation. We previously identified a population of IL-10-producing, T follicular helper-like cells (“**Tfh10**”), linked to suppressed vaccine responses in aged mice. Here, we integrate single-cell (**sc**)RNA-seq, scATAC-seq and genome-scale modeling to characterize Tfh10 – and the full CD4^+^ memory T cell (**CD4^+^TM**) compartment – in young and old mice. We identified 13 CD4^+^TM populations, which we validated through cross-comparison to prior scRNA-seq studies. We built gene regulatory networks (**GRNs**) that predict transcription-factor control of gene expression in each T-cell population and how these circuits change with age. Through integration with pan-cell aging atlases, we identified intercellular-signaling networks driving age-dependent changes in CD4^+^TM. Our atlas of finely resolved CD4^+^TM subsets, GRNs and cell-cell communication networks is a comprehensive resource of predicted regulatory mechanisms operative in memory T cells, presenting new opportunities to improve immune responses in the elderly.

## INTRODUCTION

Aging dramatically alters immune responses. First, there are major shifts in adaptive immune cell composition that occur with age. These include a progressive loss of naïve B and T cells and an accumulation of memory T cells over time. This is due to a decline in naïve T cell output with age and intermittent encounters with pathogens and vaccines that promote the conversion of naïve T cells to effector and memory cells. Second, aging promotes the functional decline of adaptive immune cells, due to intrinsic defects within T and B cells as well as accumulation of immune-suppressive populations of T cells (Mogilenko et al. 2022; Goronzy and Weyand 2017). These immune impacts have devastating health consequences for the elderly. While many critical immune responses are muted (defense against pathogens and cancers, vaccine responses), chronic inflammation, or “inflammaging”, is also a hallmark of age-associated diseases. These complex aging phenotypes have been associated with CD4^+^ T cells, central mediators of adaptive immunity (Almanan et al. 2018; Goronzy and Weyand 2017; Kumar et al. 2018).

Single-cell genomics studies have revealed age-associated CD4^+^ T cell populations and gene signatures. For example, single-cell gene expression (**scRNA-seq**) profiling of peripheral blood mononuclear cells (**PBMCs**) from supercentenarians revealed an expanded population of CD4^+^ cytotoxic T lymphocytes (**CTL**), whose immune defense capabilities might promote extreme longevity and healthy aging (Hashimoto et al. 2019). The largest scRNA-seq study to profile the aging CD4^+^ T cell compartment identified three age-expanded populations: CD4^+^ CTL, activated Treg subsets and a population expressing markers of exhaustion (Elyahu et al. 2019). One scRNA-seq study directly linked age-associated CD4^+^ T cell populations to immune system dysfunction: In a model of acute kidney injury, scRNA-seq profiling identified two age-associated populations of CD153^+^PD1^+^CD4^+^ T cells that promoted expansion of age-associated B cells (**ABCs**) and led to chronic inflammation (Sato et al. 2022).

Recently, we discovered a Foxp3^-^ CD4^+^ T cell population responsible for a systemic increase in IL-10 in old mice (Almanan et al. 2020). Neutralization of the IL-10 receptor restored vaccine-driven B cell responses in aged mice, while ablation of CD4^+^ Treg cells did not, suggesting involvement of a Foxp3^-^ IL-10-producing T cell population. Because many of the Foxp3^-^ IL10^+^ cells expressed T follicular helper (**Tfh**) markers (PD1, CXCR5 and IL-21) and, like Tfh, were dependent on IL-6 and IL-21, we referred to them as “**Tfh10**”. We also found evidence of age-increased IL-10 production from human Tfh. Collectively, these data suggest novel cellular and molecular strategies to improve vaccine responses in aging. Yet many questions remain. Our characterization of Tfh10 was far from exhaustive, and the relationship between the Tfh10 population with Tfh and other T helper (**Th**) subsets is unknown. It is also unclear how our age-associated Tfh10 relate to recent reports of IL-10-producing Tfh, which arose in the context of chronic lymphocytic choriomeningitis virus (**LCMV**) infection (Xin et al. 2018) or potential regulatory control of atopy by human tonsil “IL10 TF (T-Follicular)” cells (Cañete et al. 2019; Kumar et al. 2021). Deep sc-genomics profiling of the CD4^+^ memory T cell compartment, and IL-10-producing cells in particular, would help address these critical questions.

Molecular network modeling would advance our understanding even further, elucidating mechanisms of regulatory control that could eventually be targeted to improve CD4^+^ memory T (**CD4^+^TM**) cell immune responses in aging. Specifically, gene regulatory network (**GRN**) models are critical to reprogramming cellular behavior, as they connect context-and cell-type-specific gene expression programs to transcription factor (**TF**) regulators (Bonneau et al. 2007; Davidson 2010). Reverse-engineering GRNs from gene expression data is a challenging problem, in part owing to high dimensionality. Hundreds of TFs, including those with correlated expression patterns, are detected in RNA-seq, and thus there are often many GRN models (sets of TF regulators) that are consistent with the data (i.e., could explain a given gene’s expression pattern). To improve identifiability of the correct model, TF binding predictions from chromatin accessibility data (e.g., ATAC-seq) can serve as valuable constraints for GRN modeling from gene expression data and boost GRN inference accuracy (Duren et al. 2017; Miraldi et al. 2019; Blatti et al. 2015). Importantly, both RNA-seq and ATAC-seq have minimal sample input requirements (including standard single-cell protocols), enabling application of our GRN inference method to rare cell types and *in vivo* settings. Indeed, we rigorously benchmarked our method in a well-studied *in vitro* Th17 system (Miraldi et al. 2019) before applying to intestinal innate lymphoid cells *in vivo* (Pokrovskii et al. 2019). The method was recently benchmarked on single-cell data as well (Gibbs et al., 2022). As a complement to GRNs, new methods for cell-cell communication network modeling from scRNA-seq (Efremova et al. 2020; Jin et al. 2021; Browaeys et al. 2020) can nominate extracellular signals that rewire GRNs and alter cell behavior. Together, these unbiased, genome-scale molecular network modeling techniques can delineate and prioritize regulatory mechanisms underpinning age-dependent immune dysfunction, distilling novel, actionable hypotheses and unexpected insights from sometimes overwhelmingly complex, multi-modal sc-genomics data.

Here, we use scRNA-seq and scATAC-seq to deeply profile the CD4^+^TM cell compartment, enriching for IL-10^+^ cells, to capture the “Tfh10” population we linked to suppressed vaccine responses in aged mice. We identified 13 populations of CD4^+^TM cells, including two IL-10^+^ populations that expanded with age (Tfh10 and *Prdm1^+^*CD4^+^ CTL). We find support for these populations through cross-comparison to previous sc-genomics aging studies and directly compare our age-associated Tfh10 to Tfh10 identified in other contexts. Through integration with pan-cell sc-genomics studies, we nominate intercellular signaling interactions and microenvironmental cues that might drive age-dependent accrual of Tfh10 and cytotoxic CD4^+^ T cells. Finally, we integrate our scRNA-seq and scATAC-seq into GRNs, to predict age-dependent and cell-type-specific TF regulatory mechanisms. Our integrated experimental-computational modeling approach expands our understanding of aging on the TM compartment, from TF control of IL-10-producing cells to extracellular signals supporting age-dependent CD4^+^TM and their GRNs.

## RESULTS

### Aging shifts IL-10 production from Treg to Tfh10 and *Prdm1*^+^ cytotoxic CD4^+^ cells

We globally assessed the regulatory landscape of Tfh10 and the full CD4^+^ memory T (**CD4^+^TM**) cell compartment. We performed scRNA-seq and scATAC-seq on splenic CD4^+^TM cells from young (≤ 4 months) and old (≥ 18 months) C57BL/6 mice, enriching for IL-10^+^ and IL-10^-^ cells using VertX (IL-10-IRES-GFP) mice (Madan et al. 2009; Almanan et al. 2020) (**Figs. 1A, S1A**). The reporter strategy proved essential, as detection of *Il10* from scRNA-seq or accessibility of the *Il10* locus from scATAC-seq in individual IL-10^+^ cells was variable due to drop-out or undersampling (Bacher and Kendziorski 2016; Kharchenko et al. 2014) (**Fig. S1B**). Averaging across cells to create “pseudobulk” signal tracks for each sample, *Il10* expression and accessibility were higher in GFP^+^ relative to GFP^-^ cells, as expected (**Fig. S1C**, P<10^-7^).

**Figure 1:**
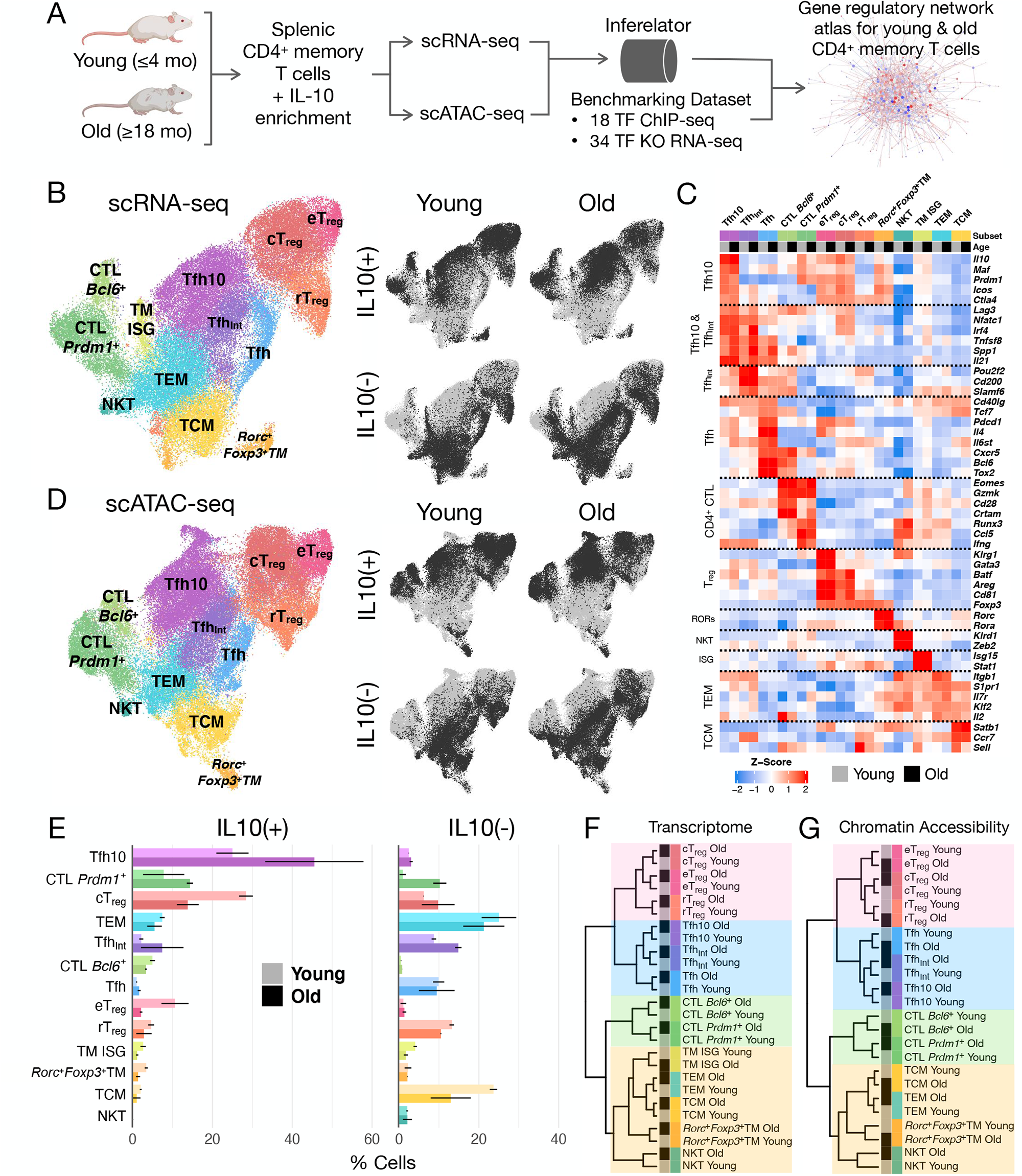
Aging shifts IL-10 production from T_reg_ to IL-10-producing Tfh-like (Tfh10) and *Prdm1*^+^ CD4^+^ cytotoxic (CTL) cells. **(A)** Study design for sc-genomic characterization and gene regulatory network (GRN) inference for splenic CD4^+^ memory T (CD4^+^TM) cells in young and old mice. **(B)** UMAP visualization of the 13 CD4^+^TM populations (left), with annotation of age group and IL-10 status highlighted in black (right). **(C)** Gene expression of select markers in each T-cell population and age group (pseudobulk signal, averaged across n=2 biological replicates). **(D)** UMAP visualization of scATAC-seq, with cell populations (left) and IL-10 and aging status (right) indicated. **(E)** The distribution of T-cell populations in the IL-10^+^ and IL-10^-^ compartments, estimated from scRNA-seq. Light and dark colored bars correspond to young and old age groups, respectively. Error bars indicate standard deviation (n=2). **(F-G)** Genome-scale, hierarchical clustering of the CD4^+^TM populations, based on pseudobulk (F) transcriptome and (G) chromatin accessibility profiles.

Clustering of the sc-transcriptomes yielded 13 populations of CD4^+^TM cells (**Fig. 1B**). To complement marker-based cell-type annotations (**Fig. 1C**), we explored cell-type relationships at genome scale. Hierarchical clustering of transcriptome and accessibility profiles revealed concordant relationships among populations (**Figs. 1F, G**, see **Methods**). By both scRNA-seq and scATAC-seq, the populations clustered into four major groups: regulatory T cells (**T_reg_**), Tfh-like, cytotoxic T lymphocytes (**CTL**) and other TM populations. Cell type, as opposed to age, was the major source of variation in both data types, as age groups within each subset clustered most similarly.

The Tfh-like group was composed of three populations: “classic” Tfh, the suspected “Tfh10” from (Almanan et al. 2020) and an intermediate Tfh population. As expected, the Tfh population expressed canonical Tfh regulators *Bcl6, Tox2, and Tcf7*, signature cytokines (*Il4*, *Il21*), as well as genes critical for migration to germinal centers (*Cxcr5*) and interactions with B cells (*Pdcd1*, *Cd40lg*) (Kroenke et al. 2012).

The Tfh10 cluster resembled the age-associated IL-10^+^ Tfh-like population we previously reported (Almanan et al. 2020). They robustly expressed some canonical Tfh genes (*Cd40lg*, *Pdcd1*, *Il4*, *Il21*), expressed *Il10* more than any other CD4^+^TM population, and accumulated with age (**Figs. 1C, E**). Notably, when compared to all other CD4^+^TM populations, genomic-scale transcriptome and chromatin accessibility analyses revealed that the Tfh10 population was closely related to the Tfh population (**Figs. 1F, G**). Yet several transcripts distinguished Tfh10 from the Tfh population. While some Tfh genes (*Bcl6*, *Tox2*, *Tcf7*, *Cxcr5*) were lowly expressed in Tfh10, Tfh10 cells had the highest expression of *Il21* and *Maf*, a regulator known to promote Tfh differentiation and drive *Il4* and *Il21* expression (Kroenke et al. 2012) (**Fig. 1C**). In addition, Tfh10 expressed markers commonly associated with other CD4^+^TM subsets, such as T_reg_ signature genes (*Ctla4, Icos*, *Il10*) and the Th1-polarizing TF *Prdm1* (encoding Blimp-1), known to suppress *Bcl6* and Tfh development (Choi et al. 2020). In line with Blimp-1 activity, Tfh10 express *Ifng*. Thus, although Tfh10 are most similar to the Tfh population, they exhibit important differences.

The “intermediate” Tfh population (**Tfh_Int_**) also increased with age (**Fig. 1E**). Tfh_Int_ express a mix of Tfh and Tfh10 markers, but also exhibit unique gene expression patterns, distinct from Tfh and Tfh10 (**Fig. 1C**). For example, they uniquely express *Ccr7*, a chemokine receptor whose down-modulation (along with up-regulation of *Cxcr5*) is required for germinal center (**GC**) entry (Haynes et al. 2007). They are also defined by high expression of *Pou2f2* (encoding Oct2), a TF that promotes Tfh differentiation via *Bcl6* and *Btla* (Stauss et al. 2016). Finally, they have high expression of immune inhibitory signals *Cd200* and *Slamf6*, commonly expressed in Tfh (Crotty 2014). Together, their gene signature suggests an early Tfh polarization state, resembling “pre-Tfh”, a population characterized in a recent aging study of murine and human Tfh (Webb et al. 2021). Like the pre-Tfh cells in the prior study, the Tfh_Int_ exhibit intermediate expression of *Cxcr5, Pdcd1*, and *Bcl6*, and high *Ccr7* expression. In dynamic T-cell activation experiments, Webb *et al*. discovered an age-associated block in pre-Tfh transition to GC-Tfh, findings consistent with the compositional increase of Tfh_Int_ observed in our aged mice (**Fig. 1E**). Although Tfh_Int_ resemble pre-Tfh, our single-cell data identified an additional Tfh-like population (Tfh10), and, in the absence of dynamic single-cell data, it is not possible to make a direct comparison to Webb *et al*. Thus, we refer to our population as Tfh_Int_, because they exhibit moderate expression of both Tfh and Tfh10 markers.

Across CD4^+^TM cells, *Bcl6* and *Prdm1* expression are inversely correlated (**Fig. S1D**), consistent with their ability to mutually repress each other’s expression (Choi et al. 2020). Similar to the Tfh, this circuit also dichotomized the CTL populations (**Figs. 1B, C**). Both CD4^+^CTL express cytotoxic TF *Eomes* and mediator *Gzmk*. The “*Prdm1*^+^CD4^+^CTL” displays Th1-like polarization, including expression of *Runx3*, *Ccl5, Nkg7*, and *Ifng*. In contrast, the “*Bcl6*^+^CD4^+^CTL” drives a cytotoxic program from a Tfh-like state: They express as much *Bcl6* and *Cxcr5* as the Tfh population and have moderate expression of *Tcf7* and *Tox2*. *Bcl6*^+^CD4^+^CTL are also distinguished by high expression of *Cd28*, a co-stimulatory receptor required for CD4^+^ T helper effector responses, including Th1 or Tfh differentiation (Linterman et al. 2014). The *Bcl6*^+^CD4^+^CTL signature gene *Crtam* is known to induce CD4^+^ cytotoxicity (Takeuchi et al. 2016).

Age-dependence and IL-10 expression patterns further distinguish the two CTL populations. *Prdm1*^+^CTL are increased with age in both the IL-10^+^ and IL-10^-^ compartments (**Fig. 1E**), with high average *Il10* expression, on par with Treg populations (**Fig. 1C**). In contrast to *Prdm1^+^*CTL, *Bcl6*^+^CTL are less prevalent, not increased with age and mainly restricted to the IL-10^+^ compartment (**Fig. 1E**). Yet, on average, *Bcl6^+^*CTL *Il10* expression is less than that of *Prdm1*^+^CTL (**Fig. 1C**). Together, these results highlight an age-dependent *Prdm1*^+^CTL population with potentially substantial immune regulatory capabilities given their cytotoxic and cytokine expression profiles.

We also identified more traditional regulatory T cell populations, characterized by expression of *Foxp3* and co-inhibitory receptor *Ctla4*: (1) “conventional” Treg (**cT_reg_**), (2) effector Treg (**eT_reg_**), and (3) resting Treg (**rT_reg_**) (**Fig. 1C**). The cTreg express many well-known Treg markers including regulators *Batf* and *Ikzf2*; receptors *Cd81*, *Izumo1r*, *Icos*, and *Pdcd1*; and repair factor *Areg*. eT_reg_ expression patterns were most similar to cT_reg_ but distinguished by expression of TF *Gata3* and receptor *Klrg1*, which correspond to a highly active, terminally differentiated immunosuppressive effector state (Wohlfert et al. 2011; Yuan et al. 2014). rT_reg_ were defined by expression of the central memory marker *Sell* (encoding CD62L). Average *Il10* expression in eT_reg_ and cT_reg_ is decreased relative to Tfh10 but on par with *Prdm1*^+^CD4^+^CTL (**Fig. 1E**). *Il10* expression is low to absent in rT_reg_.

A fourth population, “*Rorc^+^Foxp3^+^*TM”, also expresses *Foxp3* and *Ctla4*, but we refrain from referring to them as Treg. At genome-scale, their transcriptional and accessibility landscapes are more similar to central and effector memory populations than T_reg_ (**Figs. 1F, G**). *Rorc^+^Foxp3^+^*TM have high expression of the Th17 regulators *Rorc* and *Rora*. Unlike T_reg_, they also express a high level of *Il7r*, that potentially suppresses IL-2 signaling critical for T_reg_ by competing with *Il2ra* (encoding CD25) for their common subunit *Il2rg* (Waickman et al. 2020).

Our analysis uncovered four additional CD4^+^TM populations: The central memory (**TCM**) cluster was marked by high expression of *Il2, Sell, Ccr7, Il7r*, and low expression of *Itgb1*, while the effector memory (**TEM**) cluster was identified based on low expression of *Ccr7* and *Sell* and high expression of *Itgb1*, *Il7r* and *S1pr1*. We also observed a Natural Killer T (**NKT**) cell population marked by high expression of TF *Zeb2* and receptor *Klrd1* as well as a population of interferon-stimulated TM (“**TM ISG**”) marked by high expression of *Isg15* and *Stat1* (**Figs. 1B, C**). These populations mainly resided in the IL-10^-^ compartment. While TCM decreased with age, the relative proportions of TEM, NKT and TM ISG exhibited little age dependence (**Fig. 1E**).

Similarly, we identified stable clusters of CD4^+^TM based on scATAC-seq profiling, yielding a total of 12 populations (**Fig. 1D**). Using gene promoter accessibility as a proxy for gene expression (see **Methods**), we mapped these populations to those characterized by transcriptome (**Fig. 1B**). Intriguingly, by scATAC-seq, we did not recover a cluster of cells corresponding to the TM ISG population. Using label transfer of cell-type annotations from scRNA-seq to scATAC-seq, cells labeled TM ISG in the scATAC-seq data were distributed across the other 12 scATAC-seq populations, suggesting that any CD4^+^TM cell is poised by chromatin accessibility to initiate a transcriptional response to interferon (**Fig. S1E**).

Based on IL10-GFP-reporter expression, the most abundant IL-10-producing populations in young cells are cTreg (28%), Tfh10 (25%) and eTreg (11%), with smaller fractions contributed by *Prdm1*^+^ CTL (8%), TEM (7%), *Bcl6^+^*CTL (5%) and rT_reg_ (5%) (**Fig. 1E**). In old mice, the two *Prdm1*^+^ CD4^+^TM populations compose 60% of IL-10^+^ cells (Tfh10 (46%), and *Prdm1*^+^CD4^+^ CTL (14%)), while cTreg contract to 14% and eTreg become negligible (2%). Also, in old mice, Tfh_Int_ (7%) and TEM (5%) contributed appreciably to the IL-10^+^ compartment. These compositional changes were confirmed by analysis of the scATAC-seq data (**Fig. S1F**).

To further validate our cellular annotations, we compared sc-resolved pseudobulk profiles to flow-sorted, bulk RNA-seq and ATAC-seq from CD4^+^TM using Foxp3-RFP; IL-10-GFP double-reporter mice. As expected, the transcriptome and chromatin accessibility profiles of our previously defined Tfh10 (Foxp3-RFP^-^IL10-GFP^+^CXCR5^+^PD1^+^) cells correlated strongly with Tfh10 (**Figs. S1G, H**). However, in old age, this flow-sorted population also had high correlation with the CTL populations, highlighting a need for new, sc-genomics-informed sorting strategies. T_reg_ (Foxp3-RFP^+^IL10-GFP^+^TM), as expected, were most correlated with cT_reg_ and eT_reg_.

Thus, our original characterization of Tfh10 (Almanan et al. 2020) was both confirmed and refined by single-cell, multi-‘omic characterization of the IL-10^+^ TM compartment in young and old mice. As previously described, aging dramatically alters the CD4^+^TM compartment, supporting accrual of hitherto poorly characterized IL-10-producing TM cells. This initial analysis revealed that, to the resolution of our data, at least two *Prdm1*^+^ populations, Tfh10 and *Prdm1*^+^ CTL, compose the previously identified, age-expanded IL-10^+^ TM cells.

### Tfh10 are identified across aging cell atlases, senescence and other contexts

Our sc-genomics design presented a unique opportunity to resolve CD4^+^TM populations in aging. While previous aging studies profiled multiple immune cell types and tissues, we focused specifically on CD4^+^ memory T cells in the spleen, providing (1) more scRNA-seq coverage of this compartment than any previous study (**Fig. 2A**) and (2) a scATAC-seq dataset of similar depth. Thus, we report more finely resolved CD4^+^TM populations than previous aging cell atlases. Furthermore, our IL-10-enrichment strategy provided increased resolution of IL-10^+^ TM subsets. Through comparison of our CD4^+^TM populations to other studies, we set out to (1) test the reproducibility/quality of our annotations and (2) to clearly delineate how our CD4^+^TM populations relate to those described in prior aging and CD4^+^TM scRNA-seq studies (**Figs. 2, S2**).

**Figure 2:**
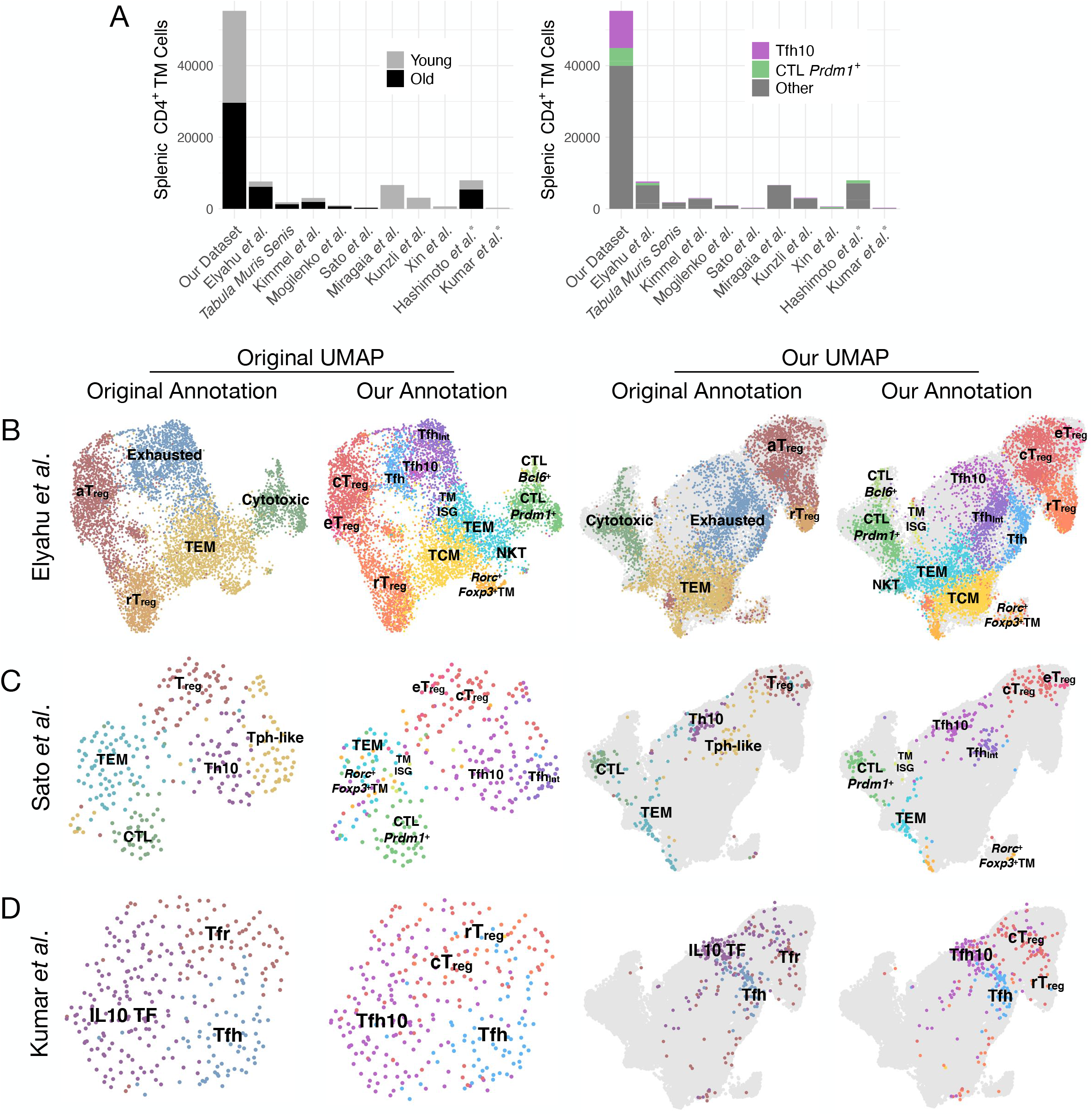
Cross-comparative analyses of other aging, Tfh10 and human cell atlases support and contextualize our highly-resolved CD4^+^TM populations. (A) Number of young and old CD4^+^ memory T cells from published scRNA-seq datasets (left) and number of Tfh10 and *Prdm1*^+^CD4^+^CTL cells identified by label transfer (right). Asterisk denotes human datasets. (B) Cells from the Elyahu *et al*. aging CD4^+^ T cell atlas projected on their UMAP coordinates (first two columns) or our UMAP coordinates (last two columns) with their original cell type annotation (first and third columns) or labels transferred from our atlas (second and fourth columns). The gray background shapes (columns 3 and 4) denote the location of cells from our atlas. (C) Senescence-associated T (SAT) cells from Sato *et al*. and (D) human tonsil Tfh, Tfr, and IL10 TF cells from Kumar *et al*. projected analogously to (B).

We first compared to (Elyahu et al. 2019), a large scRNA-seq study that characterized the full CD4^+^ T cell compartment (naïve and TM) in young (2-3 months) and old (22-24 months) mice. They identified 5 CD4^+^TM populations: cytotoxic, TEM, activated (**a**)T_reg_, resting (**r**)T_reg_ and exhausted. We applied label transfer of our cell-type annotations to their CD4^+^TM dataset. We visualized their cells, using annotations and UMAP coordinates derived from Elyahu *et al*. or our study. This side-by-side comparison of cell populations across studies provided support for our annotations (**Fig. 2B**). For example, the cytotoxic population identified in Elyahu *et al*. split cleanly into the *Prdm1*^+^ and *Bcl6*^+^ CD4^+^ CTL populations defined by our study. Similarly, their aT_reg_ split into our cT_reg_ and eT_reg_, while there was roughly one-to-one correspondence between their rT_reg_ and our rT_reg_ populations. Their “exhausted” population, classified based on *Lag3*, *Pdcd1*

(encoding PD-1), and *Tnfsf8* (encoding CD153) expression, mapped to our Tfh, Tfh_Int_ and Tfh10. We next correlated transcriptomes from CD4^+^TM in our study with those identified in Elyahu *et al*. Similarly labeled populations were highly correlated (Spearman ⍴ 0.86-0.95) (**Fig. S3A**). We also analyzed murine spleen CD4^+^TM cells from three aging cell atlases (Almanzar et al. 2020; Kimmel et al. 2019; Mogilenko et al. 2021). Despite fewer spleen CD4^+^TM in these references (**Fig. 2A**), label transfer from our reference cleanly segregated their cells in UMAP space, and we identified cells corresponding to each of our 13 CD4^+^TM populations (**Fig. S2**).

We next compared our data to a population of aging T cells arising in the context of acute kidney injury (Sato et al. 2022). In aged mice, acute kidney injury led to expansion of age-associated B cells (**ABCs**) and senescence-associated T (**SAT**) cells that interacted through CD153/CD30 signaling, to drive expansion of tertiary lymphoid tissues (**TLTs**) and chronic inflammation. SAT cells resembled Tfh10 in their production of cytokines IFNɣ, IL-21, and IL-10 and expression of old-age signature genes *Tnfsf8*, *Spp1*, and *Sostdc1*. Transferring our subset labels to scRNA-seq of aged TLT CD4^+^ T cells revealed that their two SAT populations, “Th10” and peripheral T helper-like (**Tph-like**), mapped primarily to our Tfh10 and Tfh_Int_ populations, respectively (**Fig. 2C**). In addition, their cytotoxic population aligned with our *Prdm1*^+^CD4^+^CTL. This comparison suggests that Tfh10 (1) might arise in other aging contexts (kidney injury) and (2) promote pathological B cell activation.

Next, we explored studies focused on IL-10-producing TM and/or Tfh. We examined scRNA-seq from IL-10-producing Tfh cells reported in chronic LCMV infection (Xin et al. 2018), T_reg_ and TM from spleen (Miragaia et al. 2019), and long-lived Tfh (Künzli et al. 2020). Our study suggested that IL-10 production from our age-associated Tfh10 blunted B-cell-mediated vaccine responses (Almanan et al. 2020). In contrast, IL-10 production from the Tfh10 identified by Xin *et al*. promoted humoral immunity during chronic LCMV infection (Xin et al. 2018). For scRNA-seq, Xin *et al*. used 10Bit reporter mice to isolate antigen-specific IL-10^+^CD4^+^ T cells 16 days post infection with a persistent LCMV strain. Label transfer of our annotations mapped most of their Tfh10-LCMV population to our Tfh (not Tfh10) (**Fig. S2**), despite robust detection of *Il10*. Thus, the IL-10-producing Tfh cells that arise during LCMV are both functionally and transcriptionally distinct from our Tfh10. Interestingly, other cells in their dataset (i.e., non-Tfh10-LCMV) mapped to our age-associated Tfh10. Relative to their Tfh10-LCMV, these cells expressed higher levels of several Th1-related transcripts such as *Cxcr6*, *Ccl5*, *Id2*, and *Nkg7* as well as age-associated gene *AW112010* (**Fig. S3B**). Relative to our Tfh10 cells, the Tfh10-LCMV exhibited (1) increased *Bcl6* and *Foxp3*, (2) decreased expression of TFs *Maf*, *Prdm1*, and *Jund*, and (3) decreased expression of age-associated marker *Spp1* (**Fig. S3C**).

We also compared our data to long-lived Tfh (Künzli et al. 2020). Tfh-labelled cells from Kunzli *et al*. mapped primarily to our Tfh or TCM populations. Cells mapping to Tfh10 expressed Th1 markers (e.g., *Ccl5*, *Cxcr6*, *Nkg7*, *Id2*) relative to Tfh-labeled cells in their dataset (**Fig. S2D**). Finally, as expected, we found very few age-expanded TM, including Tfh10-labeled cells, among Treg and TEM of (Miragaia et al. 2019) (**Fig. S2**).

We previously showed that aged Tfh cells from humans were enriched for IL-10 production (Almanan et al. 2020), pointing to putative Tfh10 analogs in human. To explore this possibility, we applied cross-species label transfer of our cell annotations to a scRNA-seq study of human CD4^+^TM (Kumar et al. 2021). This study characterized a subset of CD25^+^ follicular T cells in human tonsil, whose high IL-10 production dampened B cell class-switching to IgE (Cañete et al. 2019). Kumar *et al*. identified this IL-10-producing Tfh population “**IL10 TF**” from Tfh and Tfr cells isolated from human blood, lymph node, and tonsil. Their Tfh and T_reg_ populations mapped as expected to ours, and their IL10 TF population was most similar to Tfh10 (**Fig. 2D**). Examining differences in gene expression between Tfh10 and Tfh in our study and their study, our Tfh10 and Tfh cells expressed higher levels of old-age signature genes, such as *S100a10/11*, *Eea1*, and *Tnfsf8* (**Fig. S3E**).

In contrast, when we compared our data to a recent study of CD4^+^TM cells in the blood of healthy supercentenarians (Hashimoto et al., 2019), we did not identify putative Tfh, Tfh10 and several other populations (**Fig. S2**). These results highlight that tissue specificity matters when assessing CD4^+^ T cells, as some cells are present (and function) primarily within secondary lymphoid organs, while others circulate. However, we did observe that their CD4^+^ CTL population, hypothesized to promote healthy aging through immune defense, mapped to our *Prdm1*^+^ CTL cluster (**Fig. S2**).

Across aging cell atlases, the CD4^+^TM populations were recapitulated, including expansion of Tfh10 and *Prdm1*^+^CTL populations with age. This analysis linked our Tfh10 with phenotypic data in new contexts: (1) SAT Th10 that promoted ABC expansion in kidney injury (Sato et al. 2022), (2) aging spleen CD4^+^TM from later time points (Elyahu *et al*. 22-24mo, Mogilenko *et al*. 17-24mo, Kimmel *et al*. 22-23mo, *Tabula Muris Senis* 18-30mo, versus 18mo in our study). In addition, we learned that the Tfh10 associated with chronic LCMV infection (Xin et al. 2018) are both phenotypically and transcriptionally distinct from our age-associated Tfh10. Finally, we identified human populations that resemble Tfh10 (“IL10 TF” from blood, lymphoid and tonsil), providing support for human Tfh10 that complements our initial discovery of age-expanded Tfh10 in human spleen by flow cytometry (Almanan et al. 2020). We also confirmed expansion of our second age-associated IL-10-producing population, *Prdm1^+^*CTL, in other aging cell atlases, including human (Elyahu et al. 2019; Kimmel et al. 2019; Hashimoto et al. 2019).

### Resolution of age-associated gene pathways across the 13 CD4^+^ TM populations

A growing number of studies highlight age-dependent utilization of genes and gene pathways by diverse cell types, including populations composing the CD4^+^TM compartment (reviewed in (Mogilenko et al. 2022)). Here, we leverage the increased resolution of our CD4^+^TM sc-genomics atlas to pinpoint which CD4^+^TM populations exhibit specific age-dependent gene signatures, highlighting both previously characterized age-dependent pathways and those newly discovered by gene-set enrichment analysis (**GSEA**) of our data. To uncover subtle differences in our highly resolved CD4^+^TM populations, we curated custom gene sets derived from relevant T-cell gene expression studies; these complemented generic gene sets from popular databases (e.g., GO, KEGG, MSigDB; see **Methods**, **Table S1**). Our analyses focused on (1) gene pathway utilization of the 13 CD4^+^TM populations relative to each other (**Fig. 3A**, **Table S1**) and (2) identification of age-dependent gene signatures for each population (**Fig. 3B**, **Table S1**).

**Figure 3:**
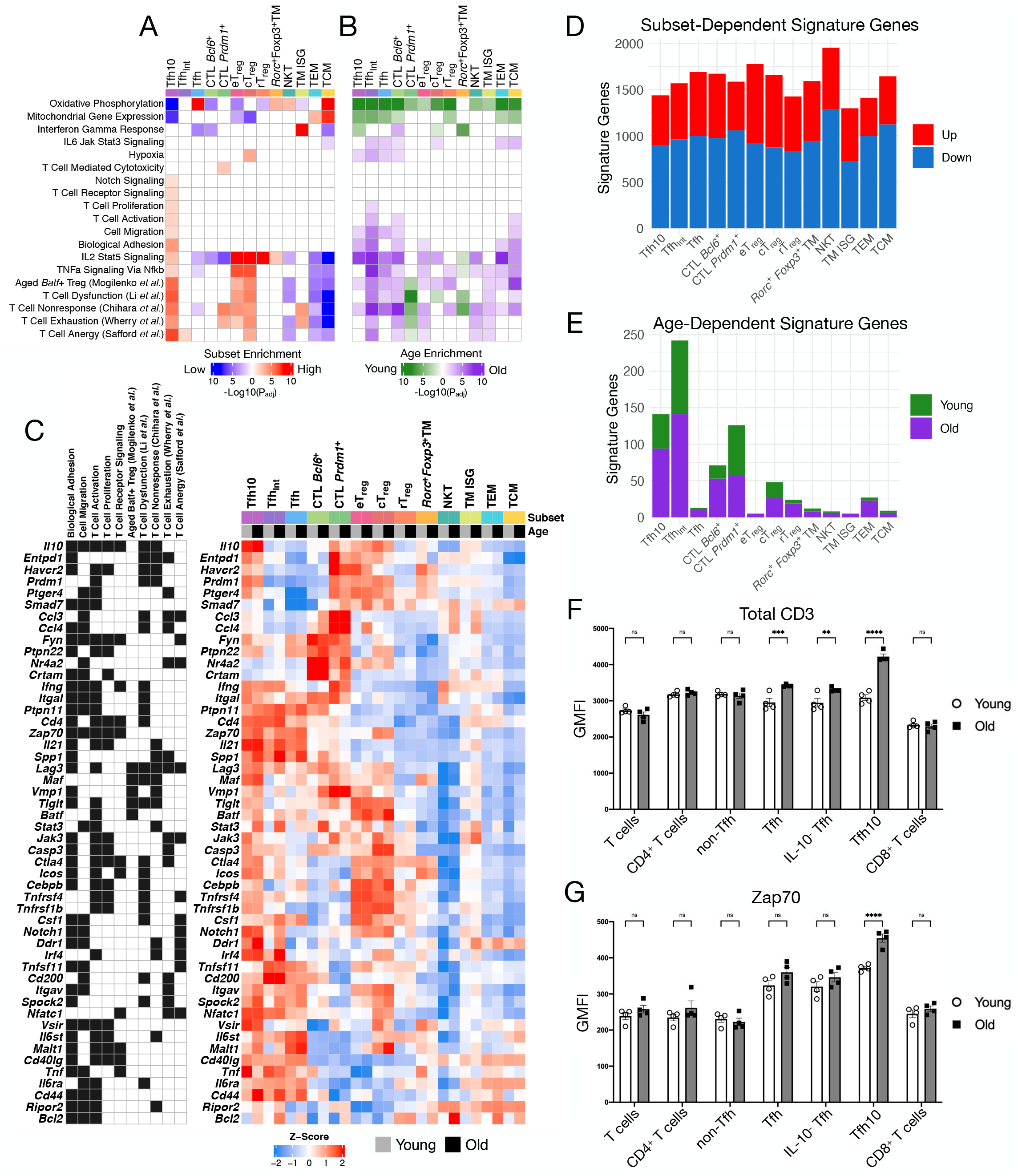
Gene pathways exhibit age-dependence and cell-type specificity within the CD4^+^TM compartment. **(A)** Gene set enrichment analysis (**GSEA**) of T cell subset-specific gene expression (select pathways shown). Red (blue) indicates high (low) enrichment (-log_10_(P_adj_), FDR=5%) of the gene pathway in the corresponding population. **(B)** GSEA of age-dependent gene expression changes. Green and purple indicate higher enrichment in young and old cells, respectively. **(C)** Genes contributing to pathway enrichments in panels (A, B) for Tfh10 and Tfh_Int_ populations related to cell adhesion, migration, and T cell-specific behavior. Left panel indicates gene pathway membership, and right panel shows gene expression in each T-cell population and age group (pseudobulk signal, averaged across n=2 biological replicates). **(D,E)** Number of (D) subset-specific and (E) age-dependent signature genes for 13 CD4+TM populations (see **Methods**). **(F,G)** Protein measurement of TCR signaling components (F) total CD3 and (G) Zap70 in splenic T cell populations from young and old mice (n=4 biological replicates). GMFI, flow cytometry geometric mean fluorescence intensity.

We identified major metabolic gene expression differences across the TM populations and with age. Across databases, the TM populations varied greatly in their utilization of oxidative phosphorylation (**ox-phos**) genes. The ox-phos pathway was strongly up-regulated in TCM and Tfh, moderately up-regulated in *Rorc^+^Foxp3*^+^TM and NKT, moderately down-regulated in *Bcl6*^+^CTL, eT_reg_, and rT_reg_, and strongly repressed in Tfh10 (**Fig. 3A**). In contrast, age-dependent utilization of the ox-phos pathway was universally decreased across TM populations, with the exception of *Rorc^+^Foxp3*^+^TM (**Fig. 3B**). Similar trends were observed for related pathways (“electron transport chain”, “cellular respiration”, “electron transfer activity”), including disease pathways with mitochondrial defects (e.g., Huntington’s disease; **Table S1**). Mitochondrial dysfunction is posited to be a major cause of maladaptive T-cell aging, impacting both memory and naïve T cells (Goronzy and Weyand 2017). Our analysis predicts that decreased mitochondrial capacity impacts nearly all CD4^+^TM populations of the aging spleen, but that the absolute levels of mitochondrial activity will vary by subset, from those with highest activity (Tfh, TCM) to lowest (Tfh10).

Based on increased expression of *Hif1a* and *Tbc1d4*, Elyahu *et al*. predicted that their age-associated “exhausted” population (which maps to our Tfh10, Tfh_Int_ and Tfh, **Fig. 2B**) was impacted by oxidative stress (Elyahu et al. 2019). Genome-scale analyses of our data support and increase the resolution of their prediction. Age-dependent expression of hypoxia genes is increased for all three Tfh-like populations, and we observe the most significant increase for Tfh_Int_. In addition, resolution of Elyahu *et al*.’s single CTL population into *Bcl6^+^* and *Prdm1^+^* populations enabled us to newly predict that the rarer “Tfh-like” *Bcl6^+^* CTL population induces hypoxia genes with age (**Fig. 3B**). Finally, while cT_reg_ expression of hypoxia genes is not age-dependent, GSEA of relative gene expression patterns across cell types revealed that hypoxia genes are uniquely up-regulated in cT_reg_, relative to the other populations. Hypoxia-induced gene expression (via hypoxia regulator Hif2a in particular) is known to support Treg homeostasis and immune suppressor function (Ben-Shoshan et al. 2008; Hsu et al. 2020).

In parallel with metabolic changes, aging for most – but not all – of the TM populations is associated with up-regulation of “T-cell dysfunction” gene signatures (**Fig. 3B**, **Table S1**). The exceptions are *Prdm1*^+^CTL and *Rorc^+^Foxp3*^+^TM, which exhibit the *opposite* trend, an age-dependent *decrease* in expression of T-cell dysfunction genes. Across TM populations, the expression of T-cell exhaustion signatures varies greatly (**Fig. 3A**, **Table S1**). These signatures are up-regulated in eT_reg_, cT_reg_, *Prdm1*^+^CTL, and Tfh10 and down-regulated in TCM, TEM and NKT. The phenotypic impact of so-called “exhaustion” signatures in Tfh10 is uncertain, given the strong overlap of exhaustion gene sets with genes important to Tfh function (*e.g.*, *Pdcd1*, *Cd200*). The exhaustion signatures overlap other gene sets, that paint a more active picture of Tfh10: T-cell activation, T-cell receptor signaling, T-cell proliferation, adhesion and migration (**Fig. 3C**). Indeed, we confirmed Tfh10 have increased TCR signaling capacity at the protein level as well (**Figs. 3F, G**). These Tfh10 gene-set associations are consistent with a senescence-associated phenotype, a supposition supported by the nearly one-to-one mapping of our Tfh10 and Tfh_Int_ to the SAT cells of (Sato et al. 2022) (**Fig. 2C**).

Of all CD4^+^TM populations, we detected the most age-dependent gene expression changes in Tfh_Int_, nearly twice as many differential genes as the next-most age-dependent populations Tfh10 and *Prdm1*^+^CTL (242, 141 and 126 differential genes, respectively, using stringent criteria (see **Methods**, **Fig. 3E**, **Table S3**)). Relative to other TM populations, Tfh_Int_ lacked unique enrichment of the gene signatures tested, as most Tfh_Int_ gene enrichments represented a diminution of either Tfh10 or Tfh enrichments (**Fig. 3A**, **Table S1**). In contrast, GSEA of age-dependent Tfh_Int_ signatures revealed dramatic shifts in Tfh_Int_ gene expression programs. Aged Tfh_Int_ adopt Tfh10 gene signatures (T cell proliferation, T cell activation, migration and adhesion, and T-cell dysfunction), suggesting that a common set of environmental signals in aging spleen might contribute to age-dependent expansion of both Tfh_Int_ and Tfh10 and/or alter the fates of Tfh_Int_, a potential source of both Tfh and Tfh10, toward Tfh10.

Having resolved gene-pathway utilizations across the CD4^+^TM (**Fig. 3**, **Table S1**), we next sought to identify the underlying gene regulatory mechanisms.

### An atlas of gene regulatory networks for TM cells in youth and old age

Gene regulatory networks (**GRNs**) describe the control of gene expression patterns by TFs. As described in the **Introduction**, integration of TF binding predictions from ATAC-seq improves GRN inference from gene expression (Miraldi et al. 2019; Pokrovskii et al. 2019; Duren et al. 2017; Blatti et al. 2015). We recently benchmarked our multi-modal modeling approach on single-cell data in yeast (Gibbs et al. 2022). Here, we undertook additional testing to ensure high-quality performance in a more complex mammalian setting. Specifically, we curated a “gold-standard” network of TF-gene interactions in CD4^+^ T cells derived from TF ChIP-seq and TF perturbation (e.g., KO) followed by RNA-seq; this network included gene targets for a total of 42 TFs (**Table S4**). We then evaluated our method for GRN inference from scRNA-seq and scATAC-seq, based on recovery of TF-gene interactions supported by TF ChIP-seq and KO (**Fig. S4**). The GRN presented here uses best practices for inference from scRNA-seq and scATAC-seq (Gibbs et al. 2022). However, for the purposes of building the highest-quality GRN for CD4^+^ T cells, we did not limit to scRNA-seq and scATAC-seq. Instead, when available, we replaced regulatory predictions from ATAC-seq with higher-quality predictions from TF ChIP-seq and TF KO (see **Methods**, **Fig. S4**), thereby leveraging valuable genomics resources generated by the community over the last 15 years.

We highlight that our GRN inference method is robust to both false positives and false negatives in prior information derived from ATAC-seq, TF ChIP-seq and TF KO, as the gene expression modeling step (1) filters TF-gene interactions in the prior not supported by the scRNA-seq data and (2) can infer new TF-gene interactions, not contained in the prior, if they are strongly supported by the gene expression model (Miraldi et al. 2019). This importantly addresses limitations of the prior information sources. For example, many of the TF ChIP-seq and TF KO come from *in vitro* contexts, while our GRN models *in vivo* gene expression of splenic CD4^+^TM. Thus, it is expected that TF-gene interactions from *in vitro* data only partially overlap with regulatory interactions occurring *in vivo*. On the other hand, although the scATAC-seq is matched to our *in vivo* context, a TF motif occurrence is not direct evidence of TF binding (introducing potential false positives) and not all TF motifs are known (introducing false negatives). Addressing these sources of error is critical to GRN inference accuracy.

#### Overview

Our GRN describes 26,745 regulatory interactions between 334 TFs and 2,675 genes. For each CD4^+^TM population, we use the GRN (knowledge of a TF’s target genes) to estimate the protein activities of TFs (**Fig. 4A**, see **Methods**). The GRN predicts subset-specific activities for many well-studied TFs. For example, Foxp3 activity is high in Treg, Bcl6 is high in Tfh, and Eomes and Runx3 are high in the CTL. However, further analysis is needed to determine whether a TF acts as an activator or repressor in a particular cell type. Thus, we identified “core” TFs, defined as TFs that contribute to the gene signature of a TM subset through (1) activation of up-regulated genes or (2) repression of genes not expressed by the subset (e.g., the signatures of other lineages). TFs acting as activators or repressors of core gene signatures are indicated in red or blue, respectively in **Fig. 4B**. Some TFs are predicted to operate as both activators and repressors, and such predictions are supported by the literature. For example, Bcl6, Prdm1 and Foxp3 are both activators and repressors of genes in Tfh, Th1 and Treg, respectively (Choi et al. 2020; van der Veeken et al. 2020). Together, these “core” analyses predict known and novel TF activities contributing to the transcriptomes, unique and shared, across the 13 CD4^+^ TM populations.

**Figure 4:**
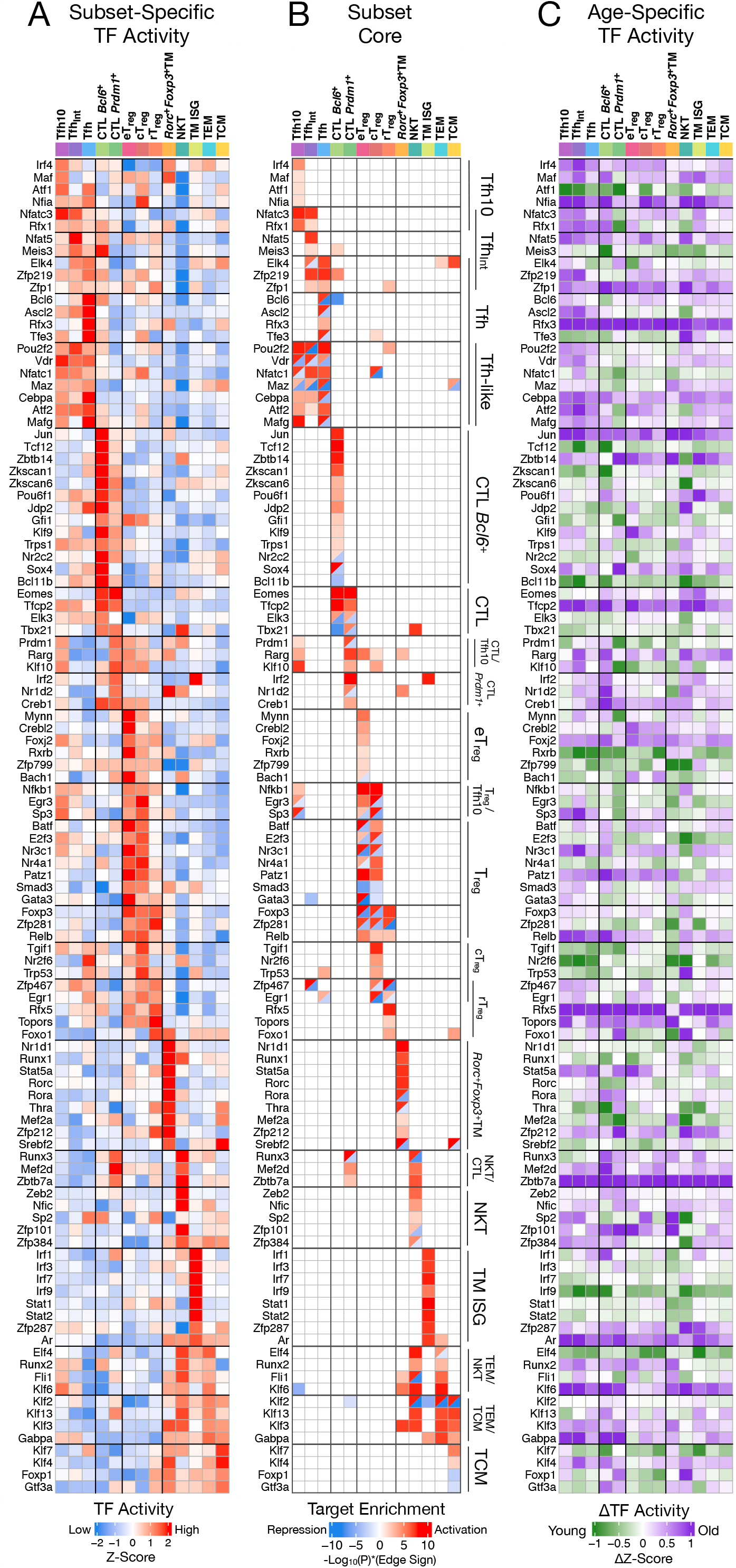
GRN modeling elucidates “core” TFs driving CD4^+^ memory T cell subset identities. **(A)** Estimated protein TF activities, averaged across age groups and biological replicates, based on the final GRN (see **Methods**, **Equation 3**). **(B)** “Core” subset-specific regulators, colored according to whether they promote subset-specific gene signatures via activation (red) or repression (blue) (see **Methods**). Red-blue triangles denote TFs that act as both activators and repressors in the given population. **(C)** Differences in TF activities between young and old populations within each T cell population. Green indicates higher activity in young, and purple indicates higher activity in old cells.

##### Age-dependent TFs

We evaluated how aging affects TF regulation in each of the subsets, calculating differential TF activity between old and young populations (**Fig. 4C**). In line with previous reports, our GRN analysis predicts that aging increases Batf activity in T_reg_ (Mogilenko et al. 2021), and, given the increased resolution of our study, we further predict that Batf activity is increased in aged Tfh_Int_, eT_reg_ and TCM populations. In line with another study characterizing aging Tfh (Webb et al. 2021), we predicted that aging promotes Irf4 and Maf activities.

However, given our comprehensive GRN modeling approach, the number of novel age-dependent TFs far exceeded those nominated by previous studies. Of the 334 TFs in our model, 166 showed age-dependent TF activity (|ΔZ-score| >1) in at least one TM population. Many of the novel TFs show stronger age dependence than those previously described. For example, Rfx3, Tfcp2 and Zbtb7a exhibit greater age-dependence than Batf, Irf4 and Maf. Furthermore, while some TF activities are concordantly age-dependent across all 13 CD4^+^TM populations (e.g., Zbtb7a), others are restricted to specific populations (e.g., increased Ascl2 activity is restricted to the aged Tfh subsets).

To help prioritize age-dependent TFs, we built models that classified young versus old cells based on TF activities (see **Methods**). These models enabled ranking of age-dependent TFs according to stability-based confidence estimates, and we highlight the 15 top-ranked age-dependent TFs (**Fig. 5**). Of the TFs with age-increased activity, several have known roles in T cell function. Regulator Stat3 has high activity in most old CD4^+^TM populations, a possible consequence of elevated, systemic IL-6 signaling in old age (Rea et al. 2018) and other age-increased signals in our dataset (*Il21*, *Il10*, *Igf1*). GRN analyses nominate Stat3 as a repressor (1) of interferon response genes (**Fig. S5**) and (2) of particular importance to old-age signatures in Tfh_Int_ and cT_reg_, as Stat3-repressed targets significantly overlap genes down-regulated in old versus young Tfh_Int_ and cT_reg_ (**Fig. S6A-C**). Zbtb7a is predicted to be a core, activating TF of *Prdm1*^+^CTL and NKT cells (**Fig. 4B**), and its age-dependent activity is highest in the old CTL populations. Zbtb7a (also known as LRF) maintains the integrity and effector potential of mature CD4^+^ T cells, partly by relieving Runx-mediated repression of *Cd4* (Vacchio et al. 2014). Thus, the age-dependent accrual of Runx3-regulated *Prdm1*^+^CTL could partially rely on this factor to maintain cytotoxic effector functions. Zfp143 is predicted to be active in nearly all old CD4^+^TM populations. This TF was previously linked to a terminally dysfunctional state of CD8^+^ T cells in the context of cancer and chronic infection (Pritykin et al. 2021). Rfx5’s activity is increased in aged Tfh/Tfh-like, CTL and Treg populations. Rfx5 is a core TF of rT_reg_ cells (**Fig. 4B**) whose predicted targets include *H2-DMa*, *H2-Oa* and *H2-Ob*; these targets are consistent with Rfx5‘s known role regulating MHC-II genes (Villard et al. 2000). Ar’s activity is highest in aged Tfh/Tfh-like and TEM/TCM populations. Ar is a core TF of TM ISG and TEM populations and a predicted positive regulator of interferon response genes (**Fig. S5**). Ar was linked to inhibition of Th1 differentiation (Kissick et al. 2014), suggesting a potential link between age-induced Ar activity and reduced Th1 differentiation in old age (Mogilenko et al. 2022). Little is known about the relationships between the remaining age-induced TFs and CD4^+^ T cell function; these include Arnt2, AU041133, Foxj3, Nfia, Zfp143, and Zfp773.

**Figure 5:**
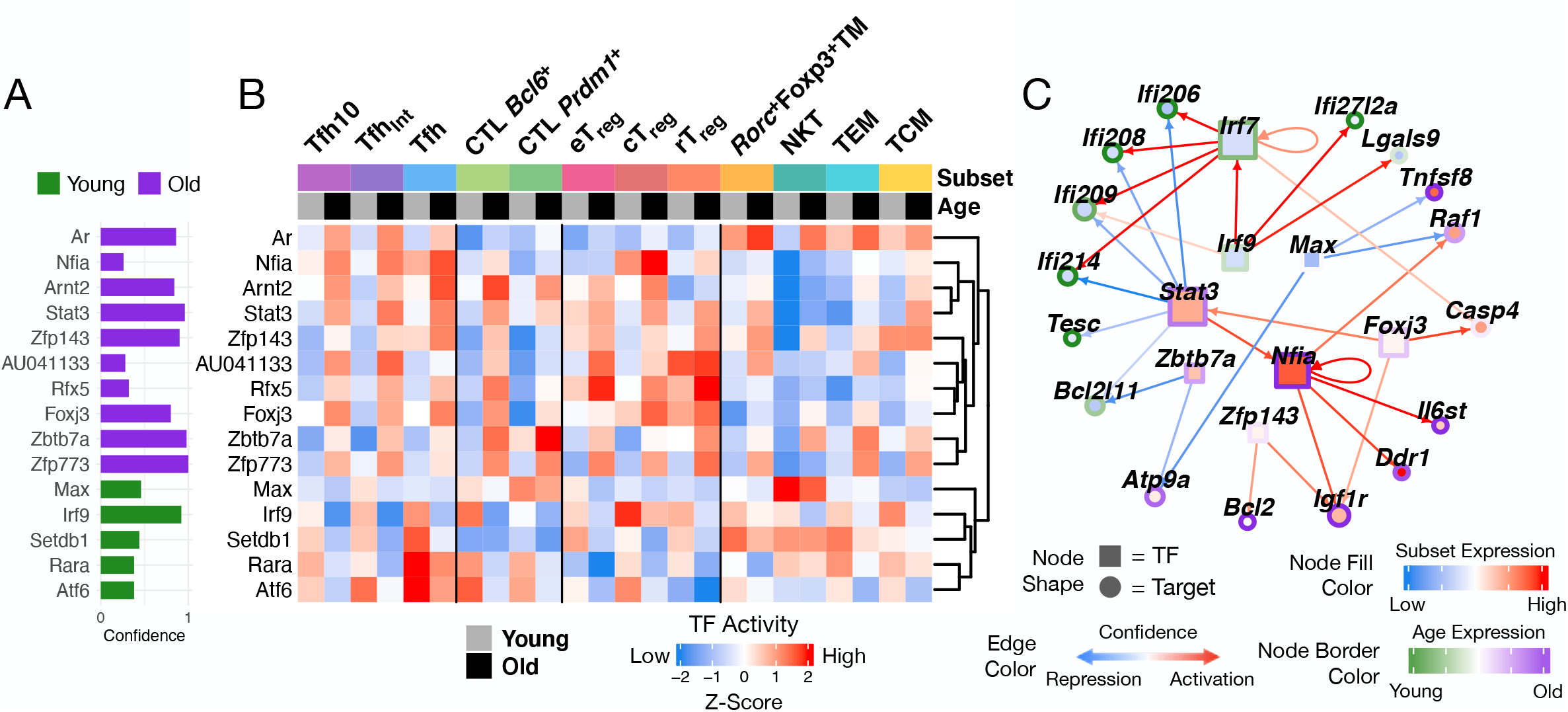
The top-ranked, age-associated TFs regulate diverse, age-associated gene pathways. **(A)** Given the large number of age-dependent TFs, we constructed a classifier to prioritize TFs whose GRN-based activities were most predictive of age. TFs were ranked according to confidence (stability across subsamples, see **Methods**). **(B)** TF activities per age group, within each T-cell population. **(C)** Select GRN interactions between age-dependent TFs and age-dependent targets within Tfh-like populations. Node fill and border colors are based on mean subset-and age-specific expression across Tfh10 and Tfh_Int_ populations, respectively. Edge color indicates either activation (red) or repression (blue) of gene expression.

Our model also prioritizes TFs with higher activities in young cells (**Figs. 5A, B**). Irf9, a well-known regulator of interferon response, has higher activity in young cells in all CD4^+^TM populations, and, as expected, its targets are enriched in interferon response genes (**Fig. S5**). Max, the obligate partner of growth-promoting TF Myc, has high activity in young *Prdm1*^+^CTL and NKT cells. Max is predicted to drive cytotoxic genes like *Fasl*, *Ctsc* and *Efhd2*. Three age-dependent TFs (Setdb1, Rara, and Atf6) have especially high activity in the young Tfh populations. Setdb1, known to inhibit Th1 differentiation (Adoue et al. 2019), is predicted to act primarily as a repressor in Tfh, controlling genes involving in T cell migration and adhesion such as *Spn*, *Cytip*, and *Ddr1*. Rara, a promoter of T cell activation (Hall et al. 2011), is predicted to drive expression of Tfh core regulator *Ascl2* and receptor *Cxcr5*. Atf6, a regulator of ER stress response (Kemp and Poe 2019), is predicted to repress *Ccdc88c*, an inhibitor of Wnt signaling that is uniquely down-regulated in Tfh. Atf6 also drives expression of mitochondria-localized *Fam162a*, contributing to the metabolic signature of Tfh.

The GRN elucidates the regulatory roles age-dependent TFs play in specific CD4^+^TM populations. For example, we visualized the regulation of select age-dependent genes in Tfh-like populations (**Fig. 5C**). The expression of interferon-stimulated genes (e.g., *Ifi206*, *Ifi208*, *ifi209*, *Ifi214*) is diminished with age. This shift is predicted to be driven by age-increased Stat3 activity and reduced Irf9 and Irf7 activities (**Fig. S5**). Along with Zbtb7a, Stat3 inhibits pro-apoptotic gene *Bcl2l11* (encoding Bim), suggesting a survival mechanism for aged cells. In addition, age-increased Zfp143 drives expression of Bim antagonist, anti-apoptotic *Bcl2*. Several TFs might indirectly promote Stat3 activity. Nfia, a Tfh10 core TF up-regulated by Stat3, activates expression of IL-6 receptor component *Il6st*. Multiple age-dependent TFs Nfia, Foxj3, and Zfp143 drive expression of IGF-1 receptor *Igf1r*; IGF-1 signaling is another known inducer of Stat3 activity that promotes age-related diseases (Salminen et al. 2021). In summary, this sub-network analysis uncovers several TFs with high activity in old age that might (1) suppress the IFN response through Stat3 activation and (2) promote survival of Tfh10 through promotion of anti-apoptotic gene expression.

### Core GRNs

For each CD4^+^TM population we curated high-confidence interactions between core TFs (**Fig. 4**) and cell-type- and/or age-specific signature genes (**Figs. 6, S6D**). These “core GRNs” highlight a subset of the regulatory mechanisms predicted to control CD4^+^TM during aging.

**Figure 6:**
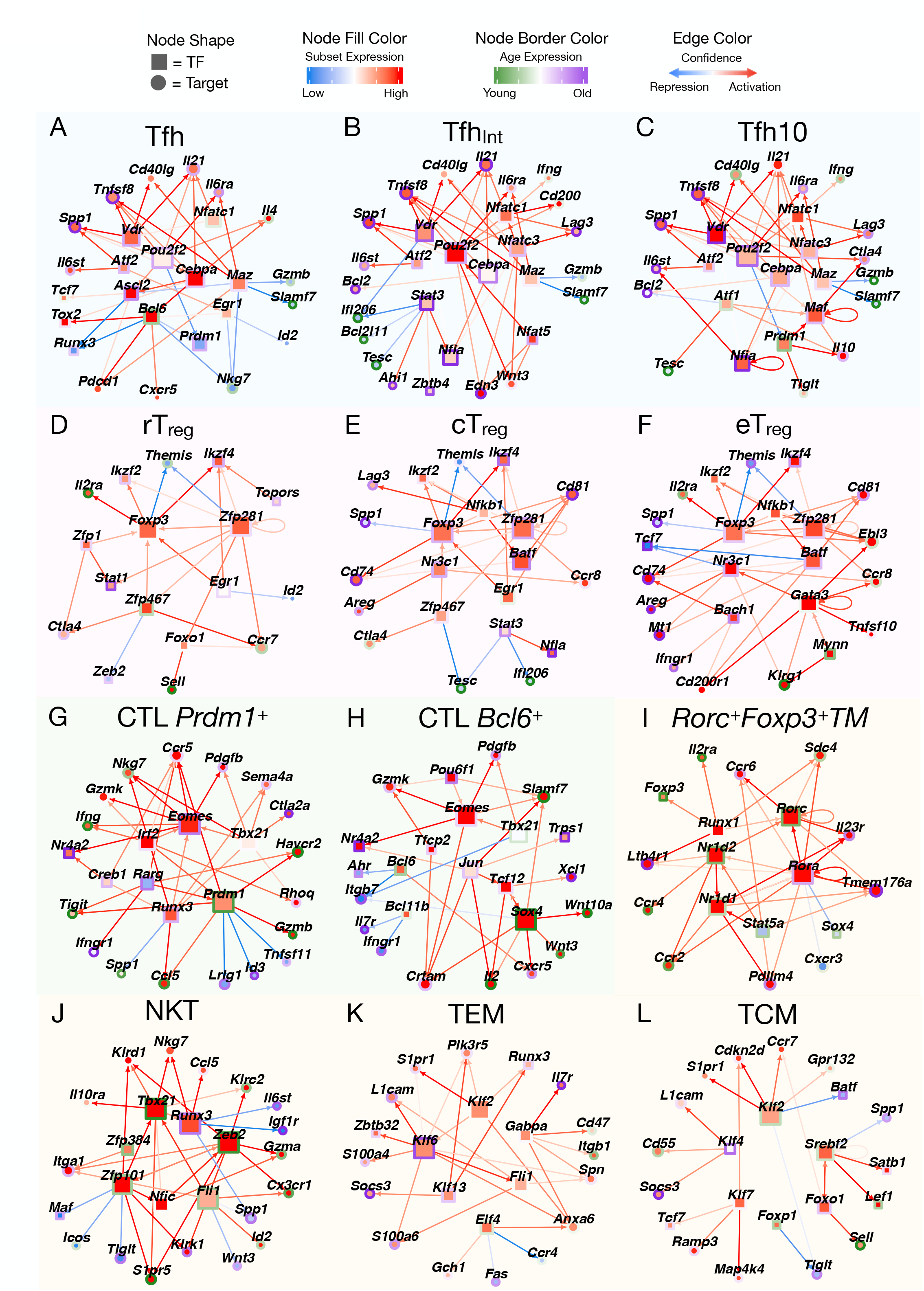
Shared and unique regulatory mechanisms drive subset- and age-specific gene expression. Selected GRN interactions between core TFs and subset- or age-specific signature genes for each T cell population (see **Fig. S6D** for TM ISG core TF network). Node shape distinguishes TFs (square) from gene targets (circle). Node fill color indicates subset-specific gene expression, and node border color indicates age-dependent gene expression within that population. Edge color indicates either activation (red) or repression (blue) of gene expression. T cell populations are grouped based on hierarchical clustering in **Fig. 1F-G**.

#### Tfh and Tfh-like populations

Core GRNs for Tfh and Tfh-like subsets (**Figs. 6A-C**) show similar regulation by densely connected TFs Cebpa, Pou2f2, Maz, Atf2 and Vdr. These TFs drive expression of genes important for Tfh differentiation and function including IL-6 receptor components *Il6ra* and *Il6st*, cytokine *Il21* and *Cd40lg*. All Tfh and Tfh-like core networks show activation of age-dependent genes *Tnfsf8* and *Spp1* (encoding osteopontin) by Cebpa, Pou2f2, Maz, Atf2 and Vdr. In Tfh_Int_ and Tfh10, NFAT factors drive age-increased expression of inhibitory receptor *Lag3*, and, in Tfh10, *Ifng* expression. These genes have been linked to Tfh-like cells that expand in old age and facilitate interactions with ABCs to promote chronic inflammation (Sato et al. 2022).

The Tfh core network (**Fig. 6A**) features the canonical Tfh regulator Bcl6 recapitulating known regulation with activation of germinal center (**GC**) Tfh markers *Cxcr5* and *Pdcd1* and repression of Th1 TFs *Prdm1* and *Runx3* (Choi et al. 2020). The Tfh core network also describes repression of Th1/cytotoxic genes including *Nkg7* (Ng et al. 2020), *Slamf7* (Cenerenti et al. 2022), *Gzmb*, and *Id2*.

Prdm1 and Maf feature prominently in the Tfh10 network. The activity of Prdm1 is increased in young cells, while the activity of Maf is increased in old. Consistent with IL-10 regulation in other CD4^+^TM populations (Zhang et al. 2020), Prdm1 and Maf are predicted to drive *Il10* expression in Tfh10. Maf also drives *Il21* (Hiramatsu et al. 2010). In addition, Prdm1 and Maf control expression of immune checkpoint inhibitory receptors *Tigit* and *Ctla4*, respectively.

Although several TM populations show age-increased Stat3 activity (**Fig. 5B**) and potential for IL-6-JAK-STAT signaling by GSEA (**Fig. 3B**), GRN analysis nominates Tfh_Int_ as the population with the highest Stat3 activity (**Fig. 5B**). Stat3’s repressed targets are highly enriched for genes down-regulated in aged versus young Tfh_Int_, highlighting an important repressor function for Stat3 in this context (**Fig. S6B**). In Tfh_Int_, Stat3 represses interferon-inducible genes (**Figs. 6B, S6B**), growth-associated *Tesc* (Kolobynina et al. 2016), and pro-apoptotic gene *Bcl2l11* (encoding Bim). The importance of Bim repression is highlighted by our prior data, showing that downregulation of Bim during aging contributes to the survival of Tfh10 cells (Almanan et al. 2020). Transcriptional control of Bim by Stat3 is supported by Stat3 knockout and ChIP-seq in IL-27-polarized naïve CD4^+^ T cells (Hirahara et al. 2015) which was incorporated into our prior-based GRN approach (see **Methods**).

GRN analysis predicts several additional mechanisms contributing to the accrual of Tfh_Int_ and Tfh10 with age (**Fig. 1E**). Expression of Bim antagonist *Bcl2* and the activity of its predicted regulator Cebpa are elevated in aged Tfh_Int_ and Tfh10, providing a second mechanism limiting apoptosis in these aged populations (**Figs. 6B, C**). In addition, we predict a mechanism of “de-repression” of anti-apoptotic *Bcl2* in aged Tfh10: The Tfh10 core TF Atf1 is predicted to be a repressor of *Bcl2*, and the activity of Atf1 is increased in young relative to old Tfh10 (**Fig. 6C**). As a last example, the IL-6/Stat3/Bim axis is reinforced through a positive feedback loop: Stat3 drives expression of age-dependent TF Nfia, a core regulator of Tfh_Int_ and Tfh10, that promotes IL-6 receptor expression (**Figs. 6B, C**). Thus, multiple age-dependent TFs (Atf1, Stat3, Nfia and Cebpa) and regulatory mechanisms contribute to an anti-apoptotic balance of *Bcl2l11* and *Bcl2*, that potentially drive accumulation of Tfh_Int_ and Tfh10 with age.

#### Treg populations

Core GRNs of T_reg_ subsets (**Figs. 6D-F**) prominently feature canonical regulator Foxp3 and TF Zfp281. These TFs, in turn, drive expression of other T_reg_-associated regulators *Ikzf2* (Helios) and *Ikzf4* (Eos) and repress *Themis*, a gene involved in T cell selection (Lesourne et al. 2009). The T_reg_ GRNs also recapitulate a known interaction between Foxp3 and *Il2ra* (encoding CD25) (Wu et al. 2006).

The cT_reg_ and eT_reg_ subsets (**Figs. 6E, F**) upregulate genes key to their regulatory functions. Nkfb1 drives inhibitory receptors *Lag3* and *Tnfrsf4* (encoding Ox40), suggesting enhanced suppressor function mediated by TNF signaling (Nagar et al. 2010). Indeed, this is consistent with GSEA, which implicated “TNFa signaling via Nfkb” in cT_reg_ and eTreg (**Fig. 3A**). In both subsets, Zfp281 and Nr3c1 drive *Ccr8*, a marker of highly suppressive T_reg_ (Whiteside et al. 2021). Nr3c1 also activates expression of repair gene *Areg*. Foxp3, Zfp281 and Batf upregulate age-increased *Cd81*, a receptor important for Treg antitumor responses (Vences-Catalán et al. 2016). Similar to the Tfh_Int_ population, cT_reg_ have age-elevated Stat3 activity, which is predicted to suppress interferon-inducible genes and drives Tfh10 core regulator Nfia, in this context as well.

The eT_reg_ subset (**Fig. 6F**), more prominent amongst young IL-10^+^ cells, bears the hallmarks of a highly activated, immunosuppressive cell state. TFs Gata3 and Bach1 are active in this subset. Gata3 activates expression of *Tnfsf10* (encoding Trail) and IL-27 subunit *Ebi3*, reinforces *Ccr8* expression and, along with Mynn, drives expression of receptor *Klrg1*. Bach1 reinforces *Cd74* expression and, in concert with Gata3 and Batf, drives expression of Cd200 receptor *Cd200r1*.

In rT_reg_, genes associated with central memory *Ccr7* and *Sell* are activated by TFs Zfp467 and Foxo1, respectively. Zfp467 and Egr1 repress factors promoting alternative subsets, *Zeb2* (NKT core TF) and *Id2* (known to promote Th1 polarization). In aged rT_reg_, Stat1 activity is increased.

#### CTL populations

The two CD4^+^TM cytotoxic subsets share core regulators Eomes and Tbx21 and Eomes-driven expression of *Gzmk* (**Figs. 6G, H**). The subsets are distinguished by expression of TFs *Prdm1* and *Bcl6*, drivers of Th1 and Tfh polarization, respectively. In the *Prdm1*^+^ CTL subset, Eomes, along with Irf2, is predicted to drive Th1 genes *Ccr5* and *Ifng* as well as cytotoxic gene *Nkg7*. As part of this circuit, Th1 TF Runx3 reinforces *Eomes* expression. Prdm1 represses Tfh gene *Id3* and activates *Gzmb*. Prdm1 activity is higher in young cells (**Fig. 4C**) and drives expression of receptors *Tigit* and *Havcr2* (encoding Tim3), immune checkpoint markers indicative of late-stage Th1 differentiation (Anderson et al. 2016). *Prdm1*^+^CTL and Tfh10 share several core TFs: Prdm1, Klf10, and Rarg (**Fig. 4B**). Rarg shares 24/129 targets with Prdm1 (e.g., *Ccr5*, *Tigit*, *Havcr2*), but, unlike Prdm1, Rarg activity increases with age.

In the *Bcl6*^+^ CTL subset (**Fig. 6H**), Bcl6 is predicted to activate GC marker *Cxcr5* and repress T_reg_ receptor *Ahr*. Sox4 also has high activity in *Bcl6*^+^ CTL, especially in young cells, and is predicted to drive expression of Th1 receptor and exhaustion marker *Slamf7* as well as Wnt signaling genes *Wnt3* and *Wnt10a*. Jun activity (AP-1 factor subunit) is also core to this population and elevated in old cells (**Fig. 4C**). Jun shares activating targets with Sox4, including *Slamf7* and *Crtam*, an inducer of CD4^+^ cytotoxicity. It has been shown that Jun partners with Stat3 to mediate IL-6 signaling (Ginsberg et al. 2007), possibly explaining its high activity in old cells. Sox4 and Jun also both activate *Il2*, indicative of T cell activation, while Bcl11b represses *Il7r* (encoding CD127) and *Ifngr1*.

#### Other CD4^+^TM populations

The other CD4^+^TM populations include TEM, TCM, NKT, TM ISG, and *Rorc*^+^*Foxp3*^+^TM cells (**Figs. 6I-L, S6D**). Klf2 activity is a common feature of TCM, TEM, and NKT cells. In these populations, Klf2 drives expression of trafficking receptor *S1pr1* and cell cycle inhibitor *Cdkn2d*, consistent with its roles in controlling T cell motility and quiescence (Baeyens et al. 2015). Klf2 might contribute to the balance between Tfh and other CD4^+^TM populations, as induced *Klf2* deficiency in activated CD4^+^ T cells increased Tfh cell generation, while its overexpression prevented Tfh cell production (Lee et al. 2015).

#### Rorc^+^Foxp3^+^TM

Consistent with genome-scale transcriptome and accessibility comparisons **(Figs. 1F, G)**, the *Rorc^+^Foxp3*^+^TM core GRN (**Fig. 6I**) bears little resemblance to the Treg GRNs (**Figs. 6D-F**). The canonical Treg regulator Foxp3 is not core (**Figs. 4B, 6I**), as it regulates only a few *Rorc^+^Foxp3^+^*TM signature genes: *Il2ra*, *Tgfbr1*, *Il4ra*, and *Cd72*. Instead, the *Rorc^+^Foxp3*^+^TM core features well-known Th17 TFs Rorc, Rora, Nr1d1 (Rev-ErbA) and Nr1d2 (Rev-ErbB) (Miraldi et al. 2019). While we detected little *Il17f*, *Il17a* or *Il22* (Th17 cytokines) in this population, *Rorc*^+^*Foxp3*^+^TM express *Il23r*, which, consistent with the literature, is predicted to be regulated by Rorc, Rora and Nr1d1. Chemokine receptors *Ccr2* and *Ccr4* are important for Treg and Th17 function (Iellem et al. 2001; Bakos et al. 2017); we predict both are driven by Nr1d1/2 and Rora.

#### NKT

The transcription program of NKT is very similar to NK cells and shares many features with CTL. The densely connected NK maturation-associated factor Zeb2 promotes expression of cytotoxic genes *Gzma, Klrc2* and *Klrk1* and migratory genes *Itga1*, *S1pr5* and *Cx3cr1*; Zeb2 and other drivers of the cytotoxic program, ETS-family factor Fli1 and Zfp101, are more active in young cells (**Fig. 6J**). NKT cells have high activity of Th1 TF Tbx21, which is essential for healthy NK and NKT phenotypes (Townsend et al. 2004). Tbx21 is predicted to up-regulate another cytotoxic TF *Runx3* (Taniuchi 2018; Wang et al. 2018a); both TFs are core to NKT and *Prdm1*^+^ CTL (**Figs. 6G, J**). Runx3 is predicted to activate pro-inflammatory genes *Ccl5* and *Nkg7* and repress age-increased signaling components *Il6st*, *Spp1* and *Igf1r*.

#### TEM

Krüppel-like (**KLF**) TFs (Klf2, Klf6, Klf13) play a prominent role in the TEM core GRN (**Fig. 6K**). Klf2 and Klf13 upregulate *Runx3*, which participates in T effector functions (Djuretic et al. 2007; Yagi et al. 2010). Age-increased Klf6 activity is predicted to promote expression of *S100a4*, a factor important for TCR signaling and T cell motility (Brisslert et al. 2014). ETS family TFs (Gabpa, Elf4) are also prominent. Gabpa promotes memory and effector genes *Itgb1* and *Il7r*, while Elf4 represses chemokine receptor *Ccr4*. Several TEM regulators drive expression of *Anxa6*, which regulates proliferation by maintaining CD4^+^ T cell sensitivity to IL-2 signaling (Cornely et al. 2016).

#### TCM

In the TCM core GRN, Foxo1 regulates central memory marker *Sell*, and, along with Klf2, drives *Ccr7* expression (**Fig. 6L**). Klf2 and Foxo1 are known regulators of these central memory genes, which, in turn, govern T cell homing to lymphoid tissues (Carlson et al. 2006; Fabre et al. 2008; Gray et al. 2018). Foxo1 mutually activates Srebf2, a core regulator of both TCM and *Rorc*^+^*Foxp3*^+^TM. Srebf2 drives expression of *Satb1*, a TF that prevents Tfh differentiation (Chaurio et al. 2022). Srebf2 also drives *Lef1*, a TF important for establishing T_reg_ immunosuppressive function (Xing et al. 2019). Several other KLF TFs are prominent in the TCM core. Klf7 is predicted to activate *Tcf7*, a TF important for early Tfh differentiation (Choi and Crotty 2021). Klf4 upregulates *Socs3*, an inhibitor of TCR signaling, while Klf2 and Klf7 drive cell cycle inhibitor *Cdkn2d*.

#### TM ISG

An interferon-induced regulatory program is the defining feature of TM ISG cells (**Fig. S6D**). Stat1, Stat2 and several interferon response factors (Irf1, Irf3, Irf7, Irf9) are core regulators of the TM ISG population. These activities are consistent with a Type 1 IFN response (ISGF3 complex of Stat1, Stat2 and Irf9, which drives downstream Irf1), Type 2 IFN response (signaling via Stat1 homodimer), and IFN-induced via pathogen recognition receptors (e.g., via STING and/or MAV to Irf7 and Irf3) (Schneider et al. 2014). The targets of these TFs are, as expected, enriched in IFN and response to pathogen genes (**Fig. S5**). The TF Zfp287 is also core to TM ISG. Stat2 and Zfp281 activate *Usp18*, a negative regulator of interferon signaling (Malakhova et al. 2003; Zhang and Zhang 2011). Core TM ISG regulators drive expression of *Isg15* and *Ifi44*, factors that form a complex with Stat2 in response to Type I interferon (He et al. 2021), to modulate context-dependent immune responses (Mirzalieva et al. 2022). *Ifi44* is increased in aged cells, while *Isg15* is increased in young.

### Integration of age-dependent cell-cell communication and gene regulatory networks

We previously showed that Tfh cytokines IL-6 and IL-21 were required for accrual of Tfh10 (Almanan et al. 2020). Here, we take an unbiased, systems-level approach to infer the cell-cell communication networks driving gene-regulatory and compositional changes in the aging CD4^+^TM compartment.

We first integrated our high-depth CD4^+^TM scRNA-seq with aging cell atlases that broadly profiled immune populations of the spleen (Almanzar et al. 2020; Kimmel et al. 2019) (**Figs. 7A, S6E**). This effort required significant reannotation of the pan-cell atlases. For example, for the largest, *Tabula Muris Senis*, we refined a single population of B cells (22,294 cells) into seven: age-associated B cells (**ABCs**), regulatory (**Breg**), follicular (**FOB**), germinal center (**GCB**), marginal-zone (**MZB**), transitioning (**TrB**) and cycling B cells (**Fig. S6F**). We also refined annotation of the myeloid cells, newly identifying three dendritic cell (**DC**) populations: plasmacytoid (**p**)DC, conventional (**c**)DC, and mature (**m**)DC (**Fig. S6F**). This resolution was critical, because, for example, among B cells, *Cd30* was uniquely detected in ABCs, or, among DC populations, *Il6* was uniquely detected in mDCs. Thus, high-resolution clustering was key to identifying these age-dependent signals in our downstream analysis. Of equal importance, we verified that our cell-type annotations were robustly reproduced via independent analysis of the Kimmel *et al*. atlas (**Fig. S6E**). In total, our cell-cell communication analysis included 29 immune populations.

**Figure 7:**
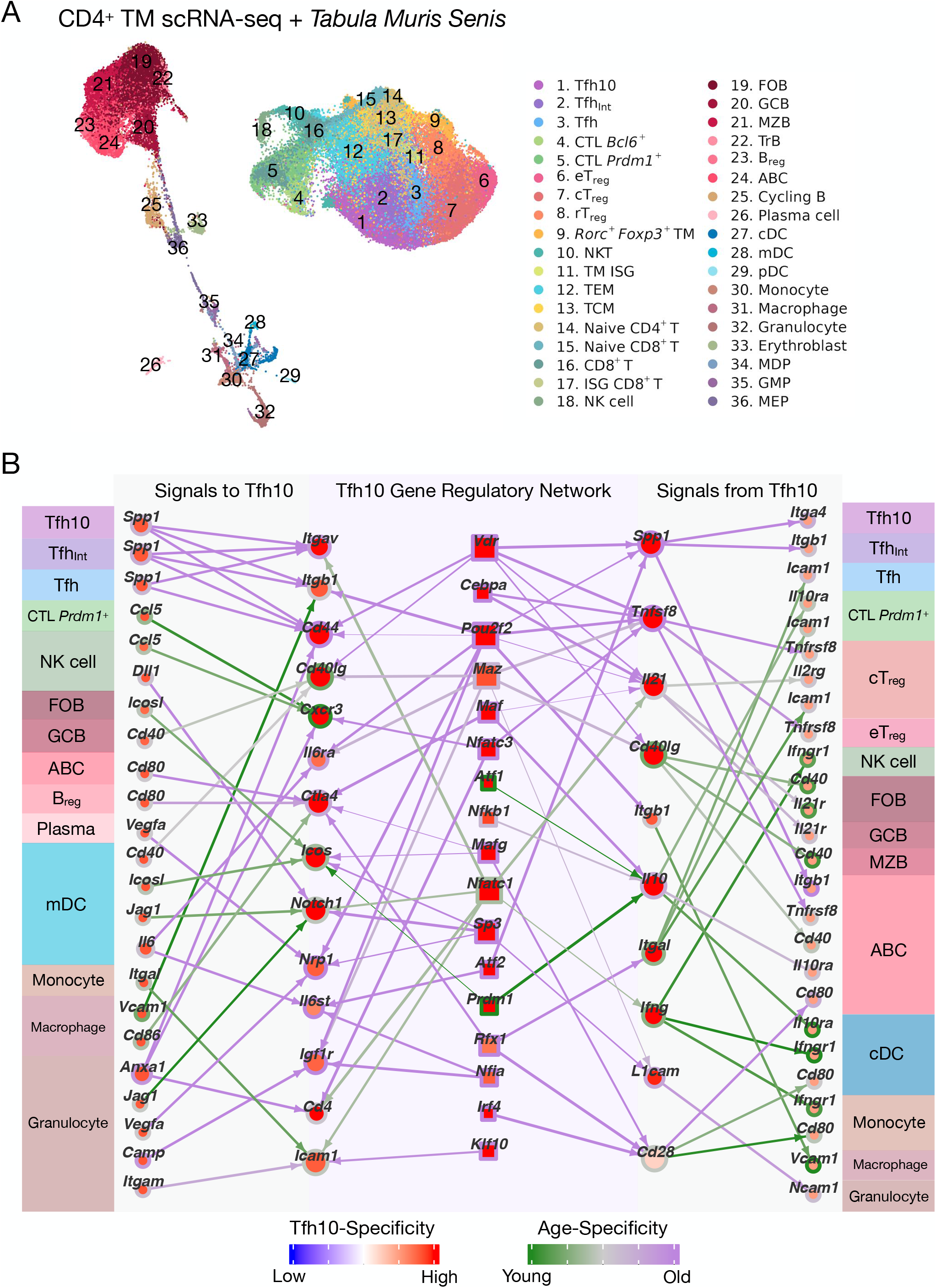
Age-dependent intercellular signals drive and are driven by Tfh10 and their GRN. **(A)** UMAP visualization of our scRNA-seq dataset integrated with age-matched, splenic cells from the *Tabula Muris Senis* atlas (Almanzar et al. 2020). **(B)** Select intercellular signals received and sent by Tfh10 along with GRN regulation of Tfh10 receptors and ligands participating in intercellular signaling. Node color of ligands (far left) and receptors (far right) are NicheNet scaled ligand and receptor scores, respectively, averaged across age (range [0,1]). Ligand and receptor border color is the difference between young and old scores in the respective cell type (range [-0.1, 0.1]). Edge color of intercellular signaling interactions is the difference in prioritization score between young and old cells (range [-0.1, 0.1]). TF (square) node color represents Z-scored TFA relative to other CD4^+^TM populations, averaged across age (range [-1, 1]). TF border and GRN interaction edge color represent difference in Z-scored TFA in old and young Tfh10 (range [-0.8,0.8]). GRN gene target (circle) node color represents Tfh10 pseudobulk expression relative to other CD4^+^TM populations, averaged across age (range [-1, 1]). GRN gene target border color represents difference in Z-scored expression between young and old Tfh10 (range [-0.8,0.8]). Abbreviations: FOB, follicular B cell; GCB, germinal center B cell; MZB, marginal zone B cell; TrB, transitioning B cell; B_reg_, regulatory B cell; ABC, age-associated B cell; cDC, conventional dendritic cell (DC); mDC, mature DC; pDC, plasmacytoid DC; MDP, macrophage-dendritic progenitor; GMP, granulocyte-macrophage progenitor; MEP, megakaryocyte-erythroid progenitor.

We performed cell-cell communication network inference using *NicheNet* (Browaeys et al. 2020). Each cell-cell signaling interaction is composed of “sender” signals (e.g., ligand) from ligand-expressing populations and “receiver” signals from cognate receptor-expressing populations. Bi-directional receptor-receptor interactions are also considered (e.g., CD40/CD40lg signaling). Receptor and ligand transcript expression is weak evidence for a signaling interaction. To improve the accuracy of such predictions, the *NicheNet* method models intracellular signaling and gene expression downstream of ligand-receptor interactions, requiring evidence of ligand-induced gene expression patterns in the “receiver” cell population. Predicted cell-cell signaling interactions, and their age-dependence, are comprehensively summarized for all cell types considered (**Fig. S7**).

At a high-level, several trends emerge (**Fig. S7**). Some of the most striking age-increased signals are predicted to impact nearly all aging immune populations. These diverse signals include Tnf super family (Tnfsf) members 4, 8, 13b and 15, apolipoprotein ApoE, Notch ligand Dll1, inducer of programmed cell death Fasl, integrin signaling and cell-cell adhesion molecules (Spp1, Fn1, Gpi1, Anxa1), chemokines Xcl1, Ccl19 and Pf4, and cytokines Il22 and Tslp. Other age-increased signals included Il21, Il4, Osm/Clcf1/Il6 (which signal to Il6ra and/or Il6st, encoding gp130), Vegfa/b and Camp (which signals to Igf1r). Several signals were predicted to be more active in young cells: chemokines Ccl3/4/5 and Ppbp, cytokines Ifng, Il2 and Tgfb1, and hormones Gcg, Adm and Gal. Other patterns include age-dependent “ligand swaps”. For example, in the case of Notch signaling, ligand Jag1 (produced by mDC and granulocytes) signals mainly to young T-cell populations, while Dll1 (produced by pDC and mDC) signals more strongly to old T-cell populations.

In **Fig. 7B**, we highlight a subset of age-dependent signaling interactions involving the Tfh10 population. We identified the cellular sources of signals to Tfh10, the cellular targets of Tfh10 signals, and the TFs predicted to regulate expression of these ligands and receptors in Tfh10. Our previous work showed that Tfh10 accrual was dependent on IL-6 and IL-21 signaling. Here, we predict mDC as a cellular source for age-increased IL-6 signaling to Tfh10 via Il6ra and Il6st. We also see evidence of age-increased Clcf1 (from all CD4+TM populations) to Tfh10 and Tfh_Int_ via Il6st. All three Tfh populations produce Il21. As expected, B-cell populations (FOB and GCB most prominently) are major targets for Il21 via Il21r, and their ligand responses are elevated in aging spleen (**Fig. S7**). Tfh10 production of signature cytokine IL-10 is predicted to impact several populations in old age: ABC, FOB and monocytes. B_reg_ and plasma cells are another source of age-increased IL-10 (**Fig. S7**. In young spleen, Tfh10, along with the CD4^+^ CTL, eT_reg_, and cT_reg_, are dominant sources of IL-10, a signal received by *Prdm1*^+^CTL, cDC and monocytes.

Our cross-study comparative analyses (**Fig. 2**) map our age-expanded Tfh10 and Tfh_Int_ populations to “SAT cells” that promoted age-dependent chronic inflammation (Sato et al. 2022). Building on this, several other signaling interactions are supported based on functional analyses of SAT. For example, the expression of *Spp1* (encoding osteopontin) was confirmed on SAT (Sato et al. 2022). Here, our analysis predicts that all Tfh populations as well as *Bcl6^+^*CTL are the producers of Spp1 in aged spleen. We further predict that Spp1 targets several integrin receptors, which predominantly impact Tfh10, Tfh_Int_ and CD4^+^ CTL via Itga4; Tfh10, Tfh_Int_, B_reg_ and ABCs via Itgb1; Tfh10, Tfh_Int_, cT_reg_, plasma B cells, monocytes and macrophage via Itgb5; or all populations via Cd44. Thus, through a variety of heterogeneously expressed receptors, our analysis highlights the age-increased, pan-cell activity of Spp1 on the full immune compartment (**Fig. S7**). In CD4^+^ T cells, Spp1 induced Th1/Th17 polarization and cytokine production (Cantor and Shinohara 2009) and might contribute to Tfh10 expression of Th1 genes (e.g., *Ifng*, *Prdm1*). Another literature-supported interaction is CD153:CD30. In our companion study, we functionally characterized the role of *Tnfsf8* (encoding CD153) in aged Tfh (Thomas et al. 2023), while a second study (Fukushima et al. 2022) showed that CD153 was critical to (1) the age-increased accrual of SAT and (2) productive TCR signaling in SAT. Here, the age-increased, bidirectional CD153:CD30 interaction is predicted to involve ABC (supported by (Fukushima et al. 2022)) and, novelly, the T_reg_ populations. CD153:CD30 interaction to T_reg_ is likely to be functionally important, too, as CD30 signaling on T_reg_ was important for their regulatory function in a model of graft-versus-host disease (Zeiser et al. 2007). Finally, Notch signaling contributed to an age-dependent defect in the differentiation of precursor Tfh to GC-Tfh, leading to an accrual of pre-Tfh that bear markers in common with our Tfh_Int_ population (Webb et al. 2021). Our analysis predicts that all T cell populations in aging spleen, including Tfh10, are exposed to age-increased Notch signaling from ligand Dll1, produced by mDC, pDC and NK cells.

Other predicted cell-cell signaling interactions involving Tfh10 are, to our knowledge, unexplored. These age-increased signals to Tfh10 include Vegfa/b (from plasma B cells and granulocytes) to Nrp1, Camp (from granulocytes) to Igf1r, and Tnfsf4 (from *Bcl6*^+^ CTL and B_reg_) to Tnfrsf4 (**Fig. 7B**). Tnfsf4 (OX40) signaling might contribute to the accrual of Tfh10, as OX40 contributed to the survival of follicular T cells and antibody responses (Gaspal et al. 2005). Three receptors are predicted to signal to Ctla4 on Tfh10: Cd80 (from Tfh, eT_reg_, ABC, B_reg_, cDC, and mDC), Cd86 (cDC, mDC and macrophage) and Icosl (from mDC). Ctla4 inhibits TCR activation, promoting anergy (Rudd et al. 2009). Ctla4 also imparts cell-extrinsic regulatory function, competing with co-stimulatory receptor Cd28 on T cells for interactions with Cd80 on APCs (Qureshi et al. 2011). These predicted Ctla4 interactions are in opposition to signals promoting TCR signaling in Tfh10 or SAT: CD30:CD153 (Fukushima et al. 2022) and Notch (Webb et al. 2021).

Some cell-cell interactions are more prominent in young Tfh10 than old; these include CCL signaling and bidirectional CAM signaling (**Fig. 7B**). Proinflammatory chemokine ligands Ccl3/4/5, expressed by a combination of *Prdm1*^+^CTL, NKT, CD8^+^ T, NK cells, and granulocytes, are predicted to target receptors Cxcr3 and Ccr5 on Tfh10. Icam1 on Tfh10 receives signals from Itgal on monocytes and Itgam on granulocytes, while Itgal1 on Tfh10 signals to Icam1 on Tfh, *Prdm1*^+^

CTL, and cT_reg_. Tfh10 is also predicted to interact with Vcam1-expressing macrophages via integrin receptor Itgb1, a costimulatory signal during CD4^+^ T cell activation (Damle and Aruffo 1991).

The Tfh10 gene regulatory subnetwork (**Fig. 7B**) highlights TFs controlling Tfh10 communication with other cell types, providing opportunities to tweak Tfh10 signaling inputs and outputs.

## DISCUSSION

The CD4^+^ memory T cell compartment is composed of functionally diverse populations, with heterogeneous impacts on the aging immune system. While some populations (e.g., CD4^+^CTL) are posited to improve immune defense capabilities and promote healthy aging, other populations (e.g., Treg, Tfh10) drive immune dysfunction, weakening defenses against pathogens and cancers and/or contributing to inflammatory states and age-associated comorbidities. Thus, specific targeting of individual CD4^+^TM subsets could provide therapeutic avenues to improve immune responses in the elderly. To potentiate this long-term goal, our study serves as an unprecedented and essential immune-engineering resource for CD4^+^TM. At the heart of our approach is integrative mathematical modeling, informed by our own scRNA-seq and scATAC-seq CD4^+^TM aging cell atlas and 133 genomics datasets from 58 independent studies.

Given the pleiotropic impacts of CD4^+^TM populations on aging immune function, single-cell resolution of the CD4^+^TM compartment is prerequisite to selectively targeting specific TM populations. From our high-depth sc-genomics data, we identified a total of 13 CD4^+^TM populations (**Fig. 1**), more than doubling the resolution of the next-largest aging CD4^+^ T cell resource (Elyahu et al. 2019). Indeed, our cell atlas neatly resolved the previously reported age-increased CD4^+^CTL, activated T_reg_ and “exhausted” populations into *Prdm1*^+^ and *Bcl6*^+^ CD4^+^CTL, cT_reg_ and eT_reg_, and Tfh, Tfh_Int_, and Tfh10, respectively. We derived independent support for our 13 CD4^+^TM populations through comparisons to previous scRNA-seq studies, spanning aging to reports of IL-10-producing Tfh in mouse and human (**Figs. 2, S2**). Based on these analyses, our Tfh10 are very likely to be SAT cells (Sato et al. 2022), connecting functional studies of SAT cells to our work in Tfh10 (Almanan et al. 2020; Thomas et al. 2023). Equally important, our Tfh10 are distinct from the similarly named “Tfh10” discovered in a chronic infection model (Xin et al. 2018). Furthermore, comparison to our bulk RNA-seq and ATAC-seq of flow-sorted Tfh10 and T_reg_ (**Figs. S1G, H**) enabled us to connect the sc-resolved populations to our previously characterized Tfh10 (Almanan et al. 2020). Refinement of cell sorting strategies, based on our atlas, will aid in the functional delineation of Tfh10 and other CD4^+^TM populations.

Having established the reproducibility of the CD4^+^TM populations, we used genome-scale modeling to elucidate the regulatory mechanisms driving their unique, shared and age-dependent transcriptomes. GSEA and cell-cell signaling network analyses (**Figs. 3, 7B, S7**) indicate that the CD4^+^TM populations have differential dependencies on gene pathways and extracellular cues. These predictions increase the resolution of previous associations for aged T cells (e.g., metabolic dysfunction, exhaustion), by pinpointing individual subsets responsible for expression of these signatures. To identify the upstream regulatory mechanisms driving these gene-expression-derived cellular “phenotypes”, we built GRN models. These models are an advance over previous CD4^+^ T cell GRN models, constructed from population-level gene expression of *in vitro* polarized or *ex vivo* sorted T cells (Ciofani et al. 2012; Gustafsson et al. 2015; Henriksson et al. 2019; Pramanik et al. 2018; Yosef et al. 2013; Miraldi et al. 2019; Zhang et al. 2020). In contrast, our CD4^+^TM GRN models describe *in vivo* regulatory mechanisms for 13 populations, resolved based on unbiased scRNA-seq. Key to our approach is integration of TF binding (inferred from our scATAC-seq or measured from ChIP-seq) and TF perturbation data, a demonstrated best practice that boosts GRN inference accuracy from gene expression data. Here, we curated a large gold-standard GRN from 15+ years of TF ChIP-seq and perturbation data, covering 42 TFs from relevant CD4^+^ T cell populations, to demonstrate that these best practices also boost GRN accuracy in a mammalian, single-cell genomics setting (**Fig. S4**). Thus, our CD4^+^TM GRN is a community resource that stands on rigorous benchmarking, rich prior genomics and novel sc-genomics data.

The CD4^+^TM GRN predicts 26,745 regulatory interactions between 334 TFs and 2,675 target genes. Our core analyses highlighted 109 TFs as “core” activators and/or repressors driving subset-specific gene signatures (**Fig. 4**), while 166 TFs show age-dependence in at least one of the 13 CD4^+^TM populations (**Figs. 4-5**). Importantly, the GRN analyses not only nominates subset-specific and age-dependent TFs but provide maps connecting them to the genes (**Figs. 5C, 6, 7B, S6D**, **Table S5**) and gene pathways they regulate (**Fig. S5**), informing how perturbation of individual TFs might impact gene expression and, by extension, function of individual CD4^+^TM populations in youth and old age. As an example among thousands, age-dependent vitamin D receptor (Vdr) is predicted to drive *Tnfsf8* (encoding CD153), *Spp1* (encoding osteopontin), and *Il21* in Tfh populations, all markers of SAT cells with CD153 being key to SAT-ABC interactions, accrual and function (Sato et al. 2022; Fukushima et al. 2022). Our cell-cell communication network predicted an additional 284 extracellular signaling interactions that are age-dependent in at least one CD4^+^TM subset (**Figs. 7B, S7**). Thus, our models provide highly-resolved, specific regulatory predictions, many of which are predicted to contribute to age-dependent immune phenotypes. Although we have integrated additional data, benchmarked and identified literature support for a subset of these interactions, the vast majority of predictions remains untested. Future experimental follow-up will be critical.

For example, our integrative computational analyses shed light on our previously reported age-dependent Tfh10 (Almanan et al. 2020), yet important questions remain. Genome-scale scRNA-seq and scATAC-seq clustering analyses clearly identify the Tfh10 cells as most similar to Tfh, providing support for our initial characterization of these cells as Tfh-like based on limited markers. Interestingly, we predict only moderate activity of the canonical Tfh regulator Bcl6 in Tfh10. Our GRN identified 21 TFs as “core” to Tfh10 function, some unique to Tfh10 (Irf4, Klf10, Maf, Rfx1, Sp3) and many shared with the other Tfh (Atf2, Cebpa, Mafg, Maz, Nfatc1, Nfatc3, Pou2f2, Vdr), T_reg_ (Egr3, Gata3, Nfkb1, Sp3, Zfp467), and *Prdm1*^+^CTL (Klf10, Prdm1, Rarg) populations (**Fig. 4B**). While gene targets downstream of these TFs provide clues about Tfh10 function (**Figs. 6C, 7B**), further functional work is critical. Our companion study provides functional evidence for a connection between one of these TFs, Maf, with IL-6 and CD153 in aged Tfh (Thomas et al. 2023). Unexpectedly, we discovered that blocking CD153 diminished vaccine responses in aged mice, suggesting that, in some aging contexts, Tfh10 might promote beneficial immune functions (Thomas et al. 2023). This recent work provides nuance to previous studies (Sato et al. 2022; Fukushima et al. 2022), including our own (Almanan et al. 2020), in which Tfh10 or highly similar SAT cells promoted deleterious age-associated phenotypes, via IL-10 and CD153:CD30 signaling.

Thus, the emerging picture of Tfh10 functionality is complex, with multiple signaling “knobs”, like IL-10, CD153 and potentially many others **(Figs. 7B, S7),** controlling Tfh10-dependent phenotypes. Our intercellular signaling analysis predicts that Tfh10 communicate with age-associated B cells through several age-dependent mechanisms (e.g., IL-10, CD153:CD30, osteopontin, CD274:CD80; **Figs. 7B, S7**), possibly supporting germinal center reactions, as in (Sato et al. 2022; Fukushima et al. 2022). The observation of our previous study that IL-10 production from a non-Treg source (which we hypothesized to be Tfh10) muted vaccine responses in old age (Almanan et al. 2020) does not necessarily conflict with the notion that Tfh10 drive B-cell responses in other contexts. The effector versus regulatory impacts of Tfh10 is a complex function, involving interactions between multiple cell types and their GRNs.

Aged Tfh10 express genes associated with exhaustion (**Fig. 3**). However, we suspect that this is due to overlap of Tfh signature genes with exhaustion signatures and that, on Tfh-like populations, so-called exhaustion genes facilitate Tfh functions. This is most strikingly supported by functional studies in SAT cells, linked to Tfh10 via transcriptome-mapping (**Fig. 2C**), in which CD153 signaling enhanced TCR signaling and proliferative capacity (Fukushima et al. 2022). Another study showed that an inflammatory, Western diet induced a population of exhausted-like, IFNɣ-expressing CD4^+^Nrp1^+^Foxp3^-^T memory cells but with high proliferative potential and capacity to migrate (Gaddis et al. 2019). Albeit a limited set of markers defining this population, they share Tfh10 genes and exhibit a phenotype (migration) consistent with GSEA predictions for Tfh10 (**Fig 3**). At a minimum, this second study provides another example of a cell population exhibiting exhaustion gene signatures but not an exhaustion phenotype.

The relationship of Tfh_Int_ to Tfh and Tfh10 is unknown. Based on limited markers, the novel Tfh_Int_ population resembles precursor Tfh discovered in mouse and humans (Webb et al. 2021). In old age, Tfh_Int_ take on Tfh10 gene signatures, suggesting that aged Tfh_Int_, a potential source of both Tfh and Tfh10, might skew toward Tfh10 (**Fig. 3A**, **Table S1**). If Tfh_Int_ are a precursor population for Tfh and Tfh10, Tfh_Int_ might be the relevant cellular target to manipulate the balance of Tfh and Tfh10 in old age. Dynamic experiments are needed to resolve these relationships.

In summary, our study resolved 13 CD4^+^TM populations in aging spleen, providing detailed maps of their GRNs in youth and old age. Throughout our study, Tfh10 serve as a motivating example for the types of predictions and insights gained from our genome-scale models. Our GRN and cell-cell signaling analyses provide an equally rich set of predictions for the other CD4^+^TM populations, including novel or less characterized populations (e.g., Tfh_Int_, the two age-increased CD4^+^CTL populations and a *Rorc^+^Foxp3^+^*TM population, which we predict to have effector rather than regulatory function). From in-depth GRN analyses to cross-study comparisons, our predictions can help guide future functional studies. Importantly, the GRNs are contextualized by age-dependent cell-cell communication networks spanning diverse immune populations, providing extrinsic and intrinsic means to alter the function/dysfunction of the CD4^+^TM compartment in aging. We hope that these resources, models and underlying sc-genomics data, serve as a rich source of regulatory hypotheses that, long-term, inspire novel therapeutic strategies to improve immune responses in the elderly.

## METHODS

### Single-cell genomics data generation

Young (≤4 months) and old (≥18 months) IL-10-GFP-reporter (VertX) mice were aged in house. All animal protocols were reviewed and approved by the Institutional Animal Care and Use Committee at the Cincinnati Children’s Hospital Research Foundation (IACUC 2019-0049). Spleens were harvested and crushed through 100-μm filters (BD Falcon) to generate single-cell suspensions. CD4^+^ memory T cells were enriched using the negative selection magnetic-activated cell sorting CD4^+^ T cell isolation kit II (Miltenyi Biotec, San Diego, CA). Enriched cells were stained with anti-CD4, anti-CD44, and anti-CD62L antibodies (BD Biosciences, San Jose, CA) and sorted for memory CD4^+^ T cells (CD4^+^CD44^hi^CD62L^lo^) that were either GFP^+^ (IL-10^+^) or GFP^-^ (IL-10^-^) by a FACSAria flow sorter (BD Biosciences, San Jose, CA).

We performed droplet-based scRNA-seq on single-cell suspensions, loading 16,000 cells per sample into each channel of the 10x Chromium Controller using the v3 Single Cell 3ʹ Reagent Kit (10x Genomics, Pleasanton, CA). We amplified barcoded cDNA by PCR and purified using SPRI bead cleanup. We prepared gene expression libraries with 50 ng of amplified cDNA, sequencing on a NovaSeq 6000 to a mean depth of 50,000 reads per cell across all samples.

We performed droplet-based scATAC-seq on single-nuclei suspensions. We isolated nuclei from sorted cell populations by resuspending sorted cell pellets in pre-chilled lysis buffer (Tris-Hcl (10mM), NaCl (10mM), MgCl2 (3mM), Tween20 (0.1%), Nonidet P40 substitute (0.1%), Digitonin (0.01%), BSA (1%) in Nuclease-free water) for 3 minutes. Chilled wash buffer was added, mixed by pipetting, centrifuged at 500 x g, and nuclei were resuspended in Nuclei buffer (10x Genomics, Pleasanton, CA). We loaded 16,000 nuclei per sample into each channel of the 10x Chromium Controller using the Single-Cell ATAC Reagent Kit (10x Genomics, Pleasanton, CA). We isolated barcoded cDNA with Dynabeads MyOne SILANE bead cleanup mixture and SPRI bead cleanup. We sequenced cDNA libraries on a NovaSeq 6000 to a mean depth of 52,000 reads per cell across all samples.

### Population-level (bulk) RNA-seq and ATAC-seq data generation

Young (≤4 months) and old (≥18 months) IL-10-GFP x Foxp3-RFP mice were generated by crossing IL-10-reporter (VertX) mice to Foxp3-IRES-mRFP mice and aged in house. As above, spleens were harvested and crushed through 100-μm filters (BD Falcon) to generate single-cell suspensions. CD4^+^ memory T cells were enriched using the negative selection magnetic-activated cell sorting CD4^+^ T cell isolation kit II (Miltenyi Biotec, San Diego, CA). Enriched CD4^+^ T cells were stained with fluorescent antibodies against CD4, CD44, CD62L, CXCR5 and PD1 (BD Biosciences, San Jose, CA) and the following sorting strategy was used to isolate T cells into 3 groups for bulk RNA-seq and ATAC-seq: “Tfh10” (CD4^+^CD44^hi^CD62L^lo^CXCR5^+^PD1^+^GFP^+^RFP^-^), Treg (CD4^+^RFP^+^) and non-Treg-non-Tfh-IL10^-^ (CD4^+^CD44^hi^CD62L^lo^CXCR5^-^PD1^-^RFP^-^GFP^-^). RNA-seq libraries were constructed using the NEBNext Ultra II directional (dUTP) RNA-seq kit. ATAC-seq was performed using the OMNI-ATAC protocol (Corces et al. 2017). Libraries were sequenced at Novogene using NovaSeq and/or HiSeq Illumina sequencers.

### Flow cytometry data generation and analysis

Single spleen cell suspensions from young and old C57BL/6 mice were generated and stimulated with PMA (25ng/ml) and Ionomycin (0.5ug/ml). After one hour of culture, monensin and Brefeldin A (10ug/ml) were added, and cells were cultured an additional 4 hours. Cells were then stained with fluoresceinated antibodies against (TCRβ, CD4, CD8, CD3, CD3ε, CXCR5, PD1, and intracellularly for ZAP70 and IL-10) (BD Biosciences, San Jose, CA) and data acquired on a FORTESSA flow cytometer and analyzed using FlowJo Software (Ashland, OR).

### Single-cell RNA-seq processing

#### Quality control

We performed alignment, initial cell barcode filtering, and UMI counting using Cell Ranger version 3.1.0 and the 10x Genomics pre-built mm10 genome reference 2020-A. We performed all downstream quality control and analyses in R version 4.0.2, using workflows from Seurat v4.04 (Hao et al. 2021). We removed cell doublets using DoubletFinder (McGinnis et al. 2019), setting the expected doublet rate to 5%. We kept quality cell barcodes with at least one UMI count from 500 genes, greater than 1,000 total UMI counts, and less than 15% of UMIs corresponding to mitochondrial transcripts. We kept genes encoding proteins and lncRNAs on autosomal and X chromosomes detected in at least 20 cells. We removed pseudogenes as well as genes associated with ribosomal RNA contamination (*Gm42418* and *AY036118)* and hemoglobin. Initial, exploratory clustering of the combined samples revealed a large population of primarily IL-10^-^ cells expressing natural killer T (**NKT**) cell markers. Because the focus of this study was characterization of conventional CD4^+^ memory T cells, we removed these cells from downstream analyses. The resulting filtered UMI counts matrix contained 55,308 cells and 20,340 genes.

#### Sample integration and clustering

We normalized expression in each cell by total UMI counts, scaled to 10,000 total counts, and log-transformed (Seurat *NormalizeData* function). Within each biological replicate, we merged samples and selected a set of 2,000 genes with most variable expression across cells based on highest variance after variance-stabilization transformation (Seurat *FindVariableFeatures* function). We mean-centered and variance-normalized expression of each highly variable gene across cells, regressing out variation due to total UMI counts (Seurat *ScaleData* function) and performed PCA dimension reduction, keeping the top 50 principal components (Seurat *RunPCA* function). We combined biological replicates using Seurat’s anchor-based integration with the top 2,000 ranked variable genes across replicates (Stuart et al. 2019). We scaled and performed PCA dimension reduction on the integrated dataset and identified clusters using the Louvain community detection algorithm (Seurat *FindNeighbors* and *FindClusters* functions). We computed the shared nearest neighbor (**SNN**) graph for 20 nearest neighbors using the top 30 principal components and identified clusters with resolution parameter 0.6. Initial clustering revealed a population of cells with high median total UMI counts and expressing markers of multiple T cell subsets. We suspected these cells to be doublets, removed them, and repeated the sample merging, replicate integration, and clustering as described above. We performed UMAP dimension reduction on the top 30 principal components (Seurat *RunUMAP* function).

#### Differential gene expression analysis

We determined differentially expressed genes between T cell subsets using DESeq2 (version 1.30.1) (Love et al. 2014). We compared populations using “pseudobulk” expression, i.e., aggregated counts from cells in the same T-cell subset, age group, and biological replicate. Squair *et al*. showed that this pseudobulk-based approach appropriately accounts for variation between biological replicates, thereby limiting false discoveries (Squair et al. 2021). However, we also leveraged the single-cell nature of our data by filtering genes from a differential expression comparison if sporadically detected in fewer than 5% of cells of a pseudobulk population under consideration. The combination of pseudobulk differential analysis with filtering of sporadically detected genes yielded signature genes per T-cell subset (see below) that were more consistent with prior knowledge than those identified using pseudobulk differential analysis alone. We robustly normalized pseudobulk expression with DESeq2 variance-stabilization transformation (**VST**).

#### Signature gene identification

For each T-cell subset, we identified a set of “signature genes” exhibiting high or low expression relative to other subsets. We used these genes for cell-type identification and to develop T cell subset-and age-specific core gene regulatory networks. As described in (Pokrovskii et al. 2019), for a given subset, we defined an upregulated signature as genes more highly expressed (Log2(FC) > 1, FDR=10%) in that subset relative to at least one other subset and not decreased (Log2(FC) < -1, FDR=10%) relative to any other subset. Similarly, for each subset, we defined a downregulated signature as genes less expressed in that subset relative to at least one other subset and not increased relative to any other subset.

#### Gene set enrichment analysis

We performed gene set enrichment analysis (**GSEA**) with the R package *fgsea* version 1.16.0 (Korotkevich et al. 2021). We downloaded Hallmark, KEGG, and Gene Ontology gene sets from MSigDB version 7.0 (Ashburner et al. 2000; Carbon et al. 2021; Kanehisa et al. 2016; Liberzon et al. 2015; Subramanian et al. 2005). We performed GSEA on DESeq2 VST-normalized pseudobulk populations, batch-corrected with ComBat (*sva* R package version 3.38.0) (Johnson et al. 2007). For T cell population-specific enrichments, we ranked genes according to z-score; specifically, we averaged expression across replicates and age groups, then mean-centered and variance-normalized (Z-scored) the expression of each gene across populations. For age-specific enrichments, we ranked genes according to the difference in average VST-normalized gene expression between old and young groups within each T cell subset.

### Single-cell ATAC-seq processing

#### Quality control

We performed alignment, initial cell barcode filtering, identification of transposase cut sites, and initial peak calling using Cell Ranger ATAC version 2.0.0 and the 10x Genomics pre-built mm10 genome reference 2020-A. We kept quality cell barcodes with 5,000-100,000 cut sites in peaks, greater than 50% fragments in peaks, and a minimum TSS enrichment score of 3 (Seurat *TSSEnrichment* function). We kept peaks on autosomal and X chromosomes. As with the scRNA-seq, we identified and removed a population of NKT cells. After filtering, the peak-level counts matrix contained 63,228 high quality cells.

#### Sample integration, clustering and reference peak set for scATAC-seq

For sample integration and clustering, we generated a new, reference peak set using the MACS2 peak-calling algorithm (version 2.1.4). Prior to peak calling, for each sample, we modified the Cell Ranger fragment file to leverage both transposase cut sites from each fragment. We created a separate entry for each end of each fragment, resulting in a bed file of all transposase cut sites. We filtered cut sites in ENCODE blacklist regions (Amemiya et al. 2019; Dunham et al. 2012). For each age group and biological replicate, we smoothed the cut sites, extending them by +/-25bp and called an initial set of peaks (MACS2 options: *--shift -25 --extsize 50 --tsize 50 --pvalue 1E-8 --keep-dup all*). We normalized each new peak-level counts matrix using run term frequency inverse document frequency (**TF-IDF**, Seurat *RunTFIDF* function) and performed latent semantic indexing (**LSI**) dimension reduction (Seurat *RunSVD* function) on all peaks with at least one cut site in 100 cells, computing the top 50 singular values. We removed reduced dimension components having Pearson correlation greater than 0.8 with total cut sites in peaks. We broadly clustered cells (Seurat *FindNeighbors* and *FindClusters* functions, resolution parameter 0.5) and merged clusters with fewer than 200 cells with their most similar cluster according to Spearman correlation of DESeq2 VST-normalized pseudobulk accessibility. Next, we performed another round of MACS2 peak calling, generating peaks for each cluster (i.e., corresponding to T cell subsets). To generate our final reference peak set, we merged cluster-specific peaks and further merged peaks across age groups and biological replicates, keeping peaks with at least one cut site in 100 cells in each replicate. The final reference peak set contained 138,334 peaks. Similar to scRNA-seq sample integration, we merged scATAC-seq samples from the same replicate and integrated replicates using Seurat’s anchor-based method on the LSI embeddings (Seurat *IntegrateEmbeddings* function). We clustered cells on the integrated LSI embedding using the Louvain algorithm with 20 nearest neighbors and resolution parameter of 0.8.

#### Label transfer to scATAC-seq and population identification

After identifying T cell subsets in the scRNA-seq dataset, we performed Seurat label transfer to cells in the scATAC-seq dataset. For scATAC-seq samples, as a proxy for gene expression, we counted cut sites in gene promoters (2,000 bp upstream to 500 bp downstream TSS, Seurat *GeneActivity* function) and normalized promoter cut sites in each cell by total counts, scaled by median total counts across cells, and log-transformed. We followed with PCA dimension reduction, anchor identification with scRNA-seq using 30 principal components, and T cell subset label transfer (Seurat *FindTransferAnchors* and *TransferData* functions). For each scATAC-seq cluster, we used a “majority vote”, labeling all cells according to the most frequent T-cell subset label within the cluster.

### Analysis of population-level genomics data

*RNA-seq.* We aligned reads from population-level, IL-10-GFP x Foxp3-RFP reporter mice RNA-seq to mm10 using Bowtie2 (option: --very-sensitive) (Langmead and Salzberg 2012). Using SAMtools (v1.9.0) (Danecek et al. 2021), we filtered reads (option: -q 30), removed duplicates, and kept reads aligning to autosomal or X chromosomes. We generated a gene-level counts matrix for all samples using the *FeatureCounts* function from the R package *Rsubread* (Liao et al. 2019). We kept genes present in the pseudobulk scRNA-seq expression matrix (described above). The resulting population-level RNA-seq counts matrix was DESeq2 VST-normalized. Finally, we calculated Z-scored Pearson correlations between each pair of population-level experiments and pseudobulk populations in young or old cells.

*ATAC-seq*. We aligned reads from population-level, IL-10-GFP x Foxp3-RFP reporter mice ATAC-seq to mm10 using Bowtie2 (option: --very-sensitive). Using SAMtools (v1.9.0) (Danecek et al. 2021), we filtered reads (option: -q 30), removed duplicates, kept reads aligning to autosomal or X chromosomes, and removed reads aligning to ENCODE blacklist regions (Amemiya et al. 2019). We shifted aligned fragment ends by 5 bp to identify Tn5 transposase cut sites. To compare population-level and single-cell ATAC-seq, the population-level cut sites were mapped to the scATAC-seq reference peak set (described above) using the *FeatureCounts* function from *Rsubread* (Liao et al. 2019). The resulting population-level ATAC-seq peaks matrix was DESeq2 VST-normalized. We kept peaks containing at least 20 counts in both population-level and pseudobulk ATAC-seq. Finally, we calculated Z-scored Pearson correlations between each pair of population-level experiments and pseudobulk populations from young or old mice.

### Label transfer to other CD4^+^ T cell scRNA-seq datasets

We curated published scRNA-seq datasets containing CD4^+^TM cells (**Figs. 2, S2**). Where possible, we obtained processed Seurat objects from the authors. Otherwise, we downloaded processed single-cell UMI counts matrices and cell-type annotations from the NCBI Gene Expression Omnibus (**GEO**; https://www.ncbi.nlm.nih.gov/geo/). We clustered cells using the standard Seurat workflow, similar to processing our scRNA-seq (described above). We kept clusters of CD4^+^TM cells based on marker gene expression. Finally, we applied Seurat anchor-based label transfer (*FindTransferAnchors* and *TransferData* functions) from our annotated CD4^+^TM scRNA-seq, annotating query cells as the reference cell-type with the highest prediction score (**Figs. 2, S2**).

### Gold-standard ChIP-seq and RNA-seq data processing for GRN evaluation and modeling

We curated a “gold-standard” network of TF-gene interactions in CD4^+^ T cells derived from TF ChIP-seq and/or TF perturbation (e.g., KO) followed by RNA-seq; this network included gene targets for a total of 42 TFs (**Table S4**). These data were first used to evaluate the GRN (**Fig. S4A-C**), and then, based on these benchmarking results, incorporated into the final network (**Table S5**) For consistency, we processed raw sequencing files for ChIP-seq and RNA-seq experiments with in-house pipelines described below. We downloaded all fastq files from GEO.

#### TF Binding

We aligned reads to mm10 using Bowtie2 (options: --very-sensitive --maxins 2000) (Langmead and Salzberg 2012). Using SAMtools (v1.9.0) (Danecek et al. 2021), we filtered reads (options: – F 1804 -q 30), removed duplicates, and kept reads aligning to autosomal or X chromosomes. Using bamutils (NGSUtils v0.5.9) (Breese and Liu 2013), we removed reads aligning to ENCODE blacklist regions (Amemiya et al. 2019). We called peaks (FDR=5%) using MACS2 (v2.1.4) (Zhang et al. 2008). For each experiment, we defined TF-gene interaction weights based on the number of ChIP-seq peaks proximal to each gene TSS (+/-5kb) and quantile-ranked interactions (from 0 to 1), averaging rankings per TFs, for TFs with multiple experiments (Ciofani et al. 2012). Average TF-gene interactions with a ranking > 0.5 were included in our ChIP-seq gold standard. Final interaction weights were Frobenius normalized per TF.

#### TF Perturbation RNA-seq

We pseudo-aligned reads to the mm10 transcriptome (Ensembl v96) using kallisto (v0.46.0) (Bray et al. 2016) and used tximport (v1.18.0) (Soneson et al. 2015) to estimate gene-level counts from transcript abundances. We identified differentially expressed genes (|Log_2_FC| > 0.5, FDR=20%) using DESeq2 (v1.30.1) (Love et al. 2014). For each experiment, we defined a set of gene targets as those differentially expressed with interaction weights of magnitude -log_10_(P_adj_). The sign of each interaction reflected the mode of TF regulation as inferred from the change in gene expression after TF perturbation; targets whose expression decreased (increased) after TF knockout have positive (negative) sign, reflecting regulatory activation (repression). As with the TF binding network, we converted interaction weight magnitudes to rank-normalized scores, averaged scores for TFs with multiple experiments, filtered interactions with cutoff 0.5, and Frobenius-normalized final interaction weights per TF.

#### Combined TF binding and perturbation network

For TFs with both binding and perturbation experiments, we combined their networks by summing the absolute values of their ranked scores, filtering interactions with cutoff 1, and Frobenius-normalizing final interaction weights. The sign of interaction was inferred based on the TF perturbation data.

### Gene regulatory network inference

#### Gene expression matrix

We inferred GRNs using DESeq2 VST-normalized pseudobulk populations, batch-corrected with ComBat (*sva* R package version 3.38.0) (Johnson et al. 2007). The resulting gene expression matrix was composed of 52 pseudobulk populations: 13 T cell subsets each with old and young age groups from 2 biological replicates. We limited the expression matrix to genes detected in at least 5% of cells within at least one T cell subset in all biological replicates. The final gene expression matrix contained 52 pseudobulk populations and 7,689 genes (**Table S6**).

#### Selection of target genes

We inferred GRNs for 2,675 genes composed of the union of (1) T cell subset-specific signature genes (**Table S2**), (2) gene differential between old and young age groups within each T cell subset (Log2(FC) > 0.25, FDR=10%, **Table S3**). We performed all differential expression analysis on pseudobulk populations as described above.

#### Selection of potential regulators

We began with a list of 1,579 putative mouse TFs collected in (Miraldi et al. 2019). For GRN inference, we limited our list of potential regulators to the union of (1) TFs that qualified as target genes, (2) TFs with greater than median variance in prior-based transcription factor activity (see Inference framework below). Our final list of potential regulators consisted of 346 TFs.

#### Prior network construction

*scATAC prior*. Similar to (Miraldi et al. 2019), we constructed a prior network of TF-target gene interactions supported by putative TF binding sites (**TFBS**) in regions of accessible chromatin proximal to target genes (**Table S7**). We used our reference peak set (described above) for TFBS prediction in accessible chromatin by motif scanning. We downloaded mouse and human motifs from the CIS-BP motif collection version 2.00 (Weirauch et al. 2014), keeping human motifs with a mouse ortholog. We scanned peaks for motif occurrences with FIMO (Grant et al. 2011) (raw P-value < 1E-5 and first-order-Markov background model). To construct our scATAC-seq prior network, TFs were associated with target genes if a motif occurrence occurred +/-10kb of the gene body. We weighted prior interactions by the number of motif occurrences in proximal peaks, based on TFBS prediction benchmarking (Cazares et al. 2023). Interaction weights for each TF were Frobenius-normalized.

*Final prior*. The prior network used for final GRN construction was informed by benchmarking (**Fig. S4A-C**). We replaced scATAC-seq-based prior interactions with those supported by gold standard ChIP-seq and/or TF knockout RNA-seq experiments (described below), when available (**Table S7**).

#### Inference framework

We model gene expression at steady state as a sparse, multivariate linear combination of transcription factor activities (**TFAs**):

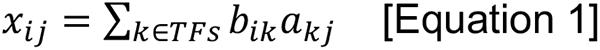

Where *x_ij_* corresponds to the expression of gene *i* in condition *j*, and *b_kj_* is the activity of TF *k* in condition *j*, and b*_ik_* describes the effect of TF *k* on gene *i*. We inferred TF-target gene interactions by solving for interaction terms b*_ik_* using the modified LASSO-StARS framework from (Miraldi et al. 2019):

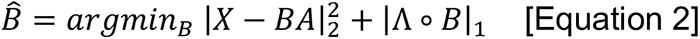

where, *X* ∈ ℝ^|*genes*|x|*samples*|^ is the gene expression matrix, *A* ∈ ℝ^|*TFs*|x|*samples*|^ specifies TF activities, *B* ∈ ℝ^|^*^genes^*^|×|^*^TFs^*^’|^ is the matrix of inferred TF-gene interaction coefficients, Λ ∈ ℝ^|genes|×|*TFS*|^ is a matrix of nonnegative penalties, and ∘ represents a Hadamard (entry-wise matrix) product. The LASSO penalty (second term in **Equation 2**) incorporates prior information into network inference. Because we seek to minimize Equation 2 we use a smaller penalty Λ*_ik_* to favor TF-gene interactions supported by the prior network (e.g., set Λ*_ik_* = 0.5 for TF-gene interactions in the prior and Λ*_ik_* = 1 otherwise). With this formulation, novel interactions can overcome the higher penalty if strongly supported by the gene expression model (first term **Equation 2**). Similar to related prior-based GRN inference methods (Greenfield et al. 2013; Siahpirani and Roy 2017; Gustafsson et al. 2015; Studham et al. 2014; Wang et al. 2022), this formulation enables: (1) refinement of the initial prior network (through removal of prior TF-gene interactions not supported by the gene expression model – i.e., robustness to false positives) and (2) the opportunity to learn new TF-gene interactions based on strong support from the gene expression data alone (i.e., robustness to false negatives). We expect both types of error in our ATAC prior, because, as examples, a motif occurrence does not necessarily correspond to TF binding (false positive) and many TF motifs are unknown, leading to false negatives. Flexible incorporation of prior information is key to accurate GRN inference.

To estimate the TF activity matrix *A*, we used two methods: (1) TF gene expression and (2) prior knowledge of TF-gene interactions, estimated from the following relationship:

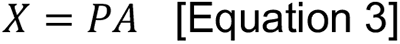

where *P* ∈ ℝ^|^*^genes^*^|×|^*^TFs^*^|^ is the prior network of TF-gene interactions. Solving **Equation 3** for *A* has improved GRN inference in diverse cell types (Miraldi et al. 2019; Pokrovskii et al. 2019; Castro et al. 2019; Tchourine et al. 2018; Arrieta-Ortiz et al. 2015; Wang et al. 2018b), including GRN inference from scRNA-seq (Jackson et al. 2020; Wang et al. 2022). However, similar to our benchmarking of GRN inference in a mammalian setting from population-level RNA-seq and ATAC-seq (Miraldi et al. 2019), modeling TF activity using TF mRNA complemented prior-based TFA in a mammalian, single-cell setting (**Fig. S4A**). Thus, we constructed GRNs using both TFA estimation methods.

For GRN model selection, we used the stability-based method StARS (Liu et al. 2010) with 50 subsamples of size 0.63 ∗ |*samples*| and an instability cutoff of 0.05 for the TF-gene interactions *B*. We ranked TF-gene interactions based on stability as described in (Miraldi et al. 2019). We generated separate GRNs using each of the TF activity estimation methods, combining predictions from the two GRNs using the maximum rank. We chose a final model confidence cutoff based on out-of-sample gene expression prediction (described below).

#### Out-of-sample gene expression prediction

As described in (Miraldi et al. 2019), we partly assessed GRN model quality and determined final model size by out-of-sample gene expression prediction. We designed two prediction tasks, inferring GRN models in the absence of (1) all young and old TEM populations (4 samples) and . (2) old rT_reg_ and cT_reg_ populations (4 samples). For each out-of-sample prediction task, we inferred GRN models using prior-based TFA and TF mRNA (see *Inference framework*). We mean-centered and variance-normalized training TFA matrices to training-set mean 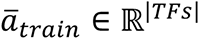 and standard deviation 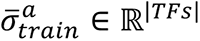. We mean centered target gene expression according to the training-set mean 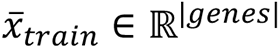. We quantified predictive performance with 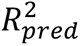 as a function of mean model size (i.e., TFs per gene). For each model-size cutoff, we regressed the vector of normalized training gene expression data onto the reduced set of normalized training TFA estimates, generating a set of multivariate linear coefficients 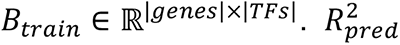 is defined as:

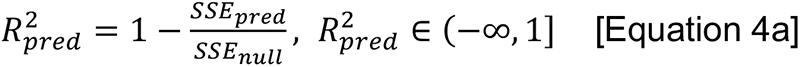

where

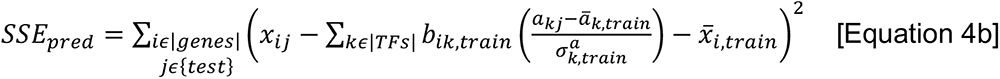

and

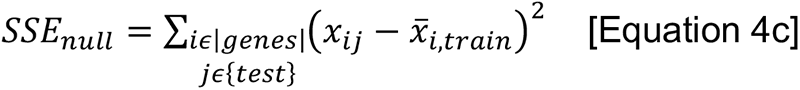

The null model used mean gene expression from the observed training data. For 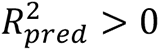 the GRN model has predictive benefit over the null model. For both out-of-sample prediction tasks, median 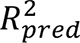 increased up to and plateaued at 10 TFs per gene (**Fig. S4D**), supporting a model-size cutoff of 10 TFs per gene.

#### Final GRN

Based on benchmarking (precision-recall analysis and out-of-sample gene expression) (**Fig. S4**), our final GRN was constructed using a prior derived from scATAC-seq, TF ChIP-seq and perturbation data (see “Prior Network Construction”) with moderate prior reinforcement (Λ*_ik_* = 0.5 for TF-gene interactions in the prior and Λ*_ik_* = 1 otherwise, see **Equation 2**). We used average network instability λ = 0.05 to rank edges by confidence. We combined GRNs based on TF mRNA and prior-based TFA by taking the maximum interaction confidence between networks, thereby preserving the individual strengths of each approach (**Fig. S4A**). We restricted the size of the final GRN to an average of 10 TFs per gene based on out-of-sample gene expression prediction (**Fig. S4D**>). The final GRN predicts 26,746 interactions between 334 TFs and 2,675 gene targets (**Table S5**).

### Gene regulatory network analysis

#### Core TF network analysis

For each T cell subset, we identified “core” TFs that promote subset-specific gene expression (**Figs. 4, 5C, 6, S6D**). Core TFs of a particular T cell subset are “activators” if their activated GRN target genes are enriched in up-regulated signature genes and “repressors” if their repressed GRN target genes are enriched in down-regulated signature genes (hypergeometric test, FDR=5%). Similarly, we identified core TFs promoting age-dependent gene expression (**Figs. S6A-C**).

#### Age-dependent TF model

We used logistic regression to build a binary classifier of age (young versus old cell populations in the CD4^+^ TM compartment) based on TF activities. TF activities were calculated from pseudobulk expression using the final GRN in **Equation 3**, as described above. Model coefficients were identified using elastic net regularization with R package *glmnet* (v4.1) (Friedman et al. 2010). The alpha parameter controlling the relative contribution of L1 to L2 penalty was set to 0.99, while we used cross validation to choose the regularization parameter lambda. Specifically, we used 50 subsamples with training set size 0.63 ∗ |*samples*| per train-test cross-validation split. We selected the regularization parameter corresponding to the sparsest model whose mean misclassification rate was within one standard error of the minimum misclassification rate. Age-dependent TFs were ranked according to a stability-based confidence: the fraction of subsamples in which a TF’s coefficient is non-zero. Age-association was based on mean partial correlation between TF activity and age across all subsamples.

#### Network visualization

We visualized GRNs using the R package *igraph* (v1.3.2) (Csardi and Nepusz 2006).

### Cell-cell signaling analysis

We refined the cell-type annotation of young and old splenic cells from the *Tabula Muris Senis* (Almanzar et al. 2020) and Kimmel *et al*. (Kimmel et al. 2019) scRNA-seq aging pan-cell atlases (**Figs. S6E, F**). First, we sub-clustered broad cell populations using the standard Seurat workflow. Then, we manually annotated sub-clusters based on expression of literature-curated panels of markers. Specifically, we resolved the T cells into naïve and memory CD4^+^ or CD8^+^ T cells and NKT cells. Memory CD4^+^ T cells were further sub-clustered and annotated by label transfer from our CD4^+^TM scRNA-seq as described above. We identified seven populations of B cells: age-associated B cells (**ABCs**), regulatory (**B_reg_**), follicular (**FOB**), germinal center (**GCB**), marginal-zone (**MZB**), transitioning (**TrB**) and cycling B cells. Finally, we refined the myeloid cells into populations of plasmacytoid (**p**)DC, conventional (**c**)DC, mature (**m**)DC, monocytes, macrophages, and granulocytes. These populations were readily identifiable in both the *Tabula Muris Senis* and Kimmel *et al*. atlases. *Tabula Muris Senis* contained some populations not identified in Kimmel *et al*. (progenitor populations, proerythroblasts, erythroblasts) possibly due to differences in protocols or count depth. These populations were not considered in the cell-cell signaling analysis.

We chose to perform the cell-cell signaling analysis using *Tabula Muris Senis* because it had greater sequencing depth. We integrated age-matched splenic scRNA-seq from *Tabula Muris Senis* with our scRNA-seq of CD4^+^TM using Seurat’s anchor-based approach with the top 2,000 ranked variable genes across datasets (**Fig. 7A**). We inferred inter-cellular signals using NicheNet (R package *nichenetr* v1.1.0) (Browaeys et al. 2020), predicting top age-specific ligand-receptor interactions between cell types. We used the differential NicheNet pipeline (github.com/saeyslab/nichenetr) with the following inputs: NicheNet’s human-to-mouse ortholog ligand-receptor network and ligand-target prior matrix, log-fold change cutoff of 0.15 for differential expression analysis, top 200 targets for calculating ligand activities, and default prioritization weights. We filtered interactions keeping the top 20 ligand signals for each cell type based on prioritization score (**Fig. S7**).

## Data Availability

Raw and processed scRNA-seq, scATAC-seq, bulk RNA-seq, and bulk ATAC-seq have been deposited in GEO under accession number GSE228668.

## ACKNOWLEDGEMENTS

We acknowledge the Cincinnati Children’s Hospital Medical Center (**CCHMC**) Gene Expression Core, DNA Sequencing Core, Flow Cytometry Core and Research IT. This work was supported by the National Institute of Health [U01AI150748 to ERM, LCK; R01AI153442 to ERM, AB; R21AI156185 to ERM; AG033057 to DAH, CAC; AG053498 to DAH, CAC].

## AUTHOR CONTRIBUTIONS

AB, CAC, DAH designed and AT, MA, MY executed the genomics and flow-cytometry experiments. JAW and ERM designed and JAW, ATB, AK and TAC executed the computational analyses. JAW and ERM led model interpretation and wrote the manuscript with input from all authors.

## DECLARATIONS

AB is a co-founder of Datirium, LLC.

## Supplemental Figure Legends

**Figure S1:**
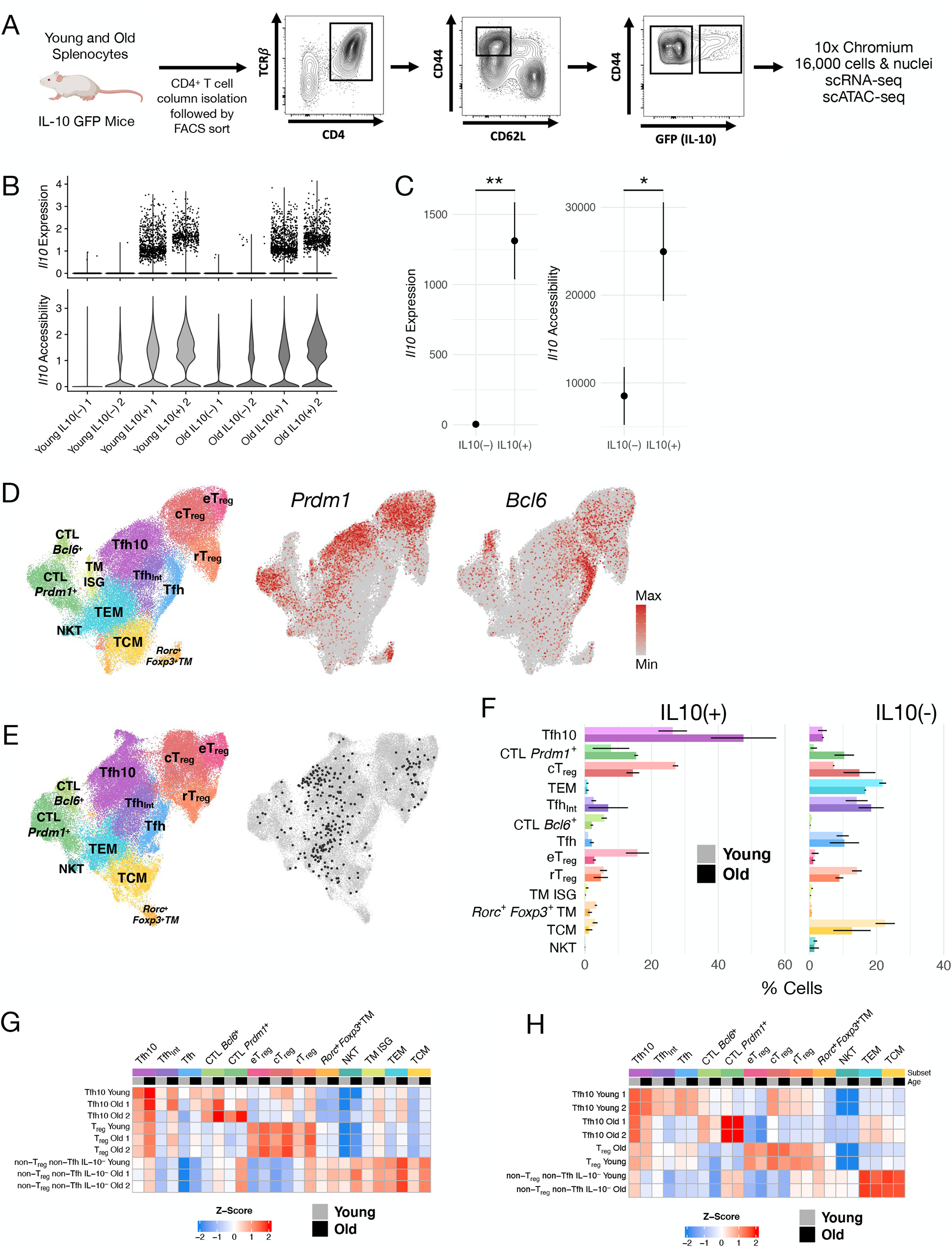
Isolation, annotation and characterization of CD4^+^ memory T cells, related to Figure 1. **(A)** Gating strategy for isolation of old and young splenic CD4^+^ memory T cells with enrichment of IL-10-GFP^+^ and IL-10-GFP^-^ cells. **(B)** Distribution of *Il10* expression (top, log-transformed transcripts per 10k transcripts) and promoter accessibility (bottom, normalized transposes cut sites within -2000 to +500bp of the *Il10* TSS) across cells per sample. **(C)** DESeq2 size factor-normalized pseudobulk *Il10* expression (left, **P<10^-74^) and accessibility (right, *P<10^-7^) for IL-10-GFP^-^ and IL-10-GFP^+^ samples. Error bars indicate standard deviation (n=2). **(D)** ScRNA-seq UMAP visualization of *Prdm1* and *Bcl6* expression. **(E)** ScATAC-seq UMAP visualization highlighting cells (black) identified by label transfer from our scRNA-seq annotation corresponding to TM ISG cells. **(F)** The distribution of T cell populations in the IL-10^+^ and IL-10^-^ compartments from scATAC-seq cells. Light and dark colored bars correspond to young and old age groups, respectively. Error bars indicate standard deviation (n=2). **(G)** Relative Pearson correlation between bulk RNA-seq and pseudobulk scRNA-seq expression. **(H)** Relative Pearson correlation between bulk ATAC-seq and pseudobulk scATAC-seq accessibility. Tfh10 (CD4^+^CD44^hi^CD62L^lo^CXCR5^+^PD1^+^GFP^+^RFP^-^), T_reg_ (CD4^+^RFP^+^) and non-T_reg_-non-Tfh-IL10^-^ (CD4^+^CD44^hi^CD62L^lo^CXCR5^-^PD1^-^RFP^-^GFP^-^) cells were sorted from IL-10-GFP x Foxp3-RFP mice.

**Figure S2:**
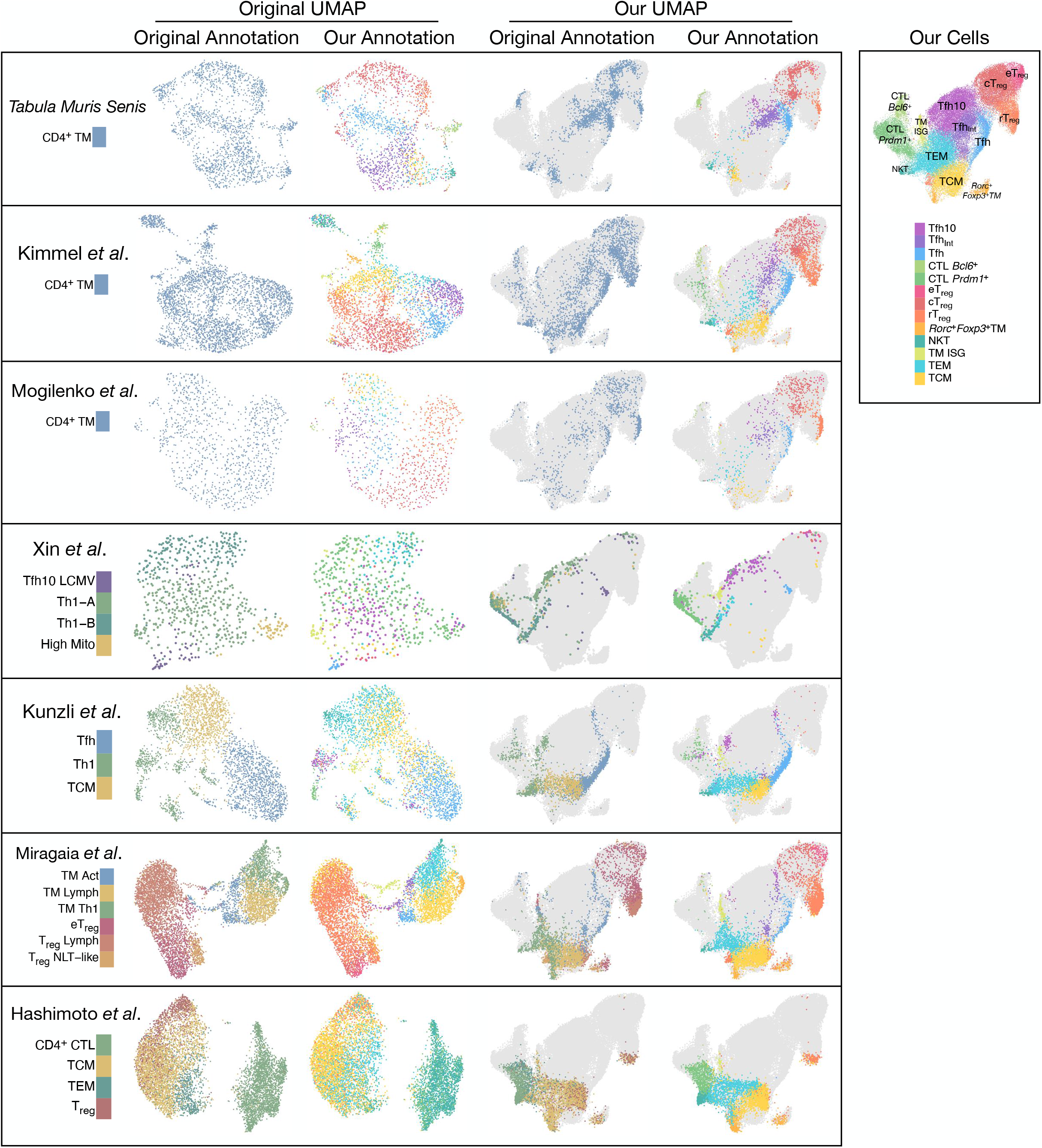
Label transfer to other CD4^+^TM scRNA-seq studies, related to Figure 2. We performed label transfer of our CD4^+^TM cell annotations to existing scRNA-seq studies of CD4^+^TM cells (Almanzar et al., 2020; Hashimoto et al., 2019; Kimmel et al., 2019; Künzli et al., 2020; Miragaia et al., 2019; Mogilenko et al., 2021; Xin et al., 2018). For each dataset, we projected cells onto UMAP coordinates from dimension reduction of the original data (first and second columns) or onto UMAP coordinates of our scRNA-seq (third and fourth columns). Gray shapes (third and fourth columns) outline the locations of cells from our scRNA-seq dataset. We annotated cells based on the original study (first and third columns, legends along left side) or based on label transfer from our scRNA-seq (second and fourth columns). Our scRNA-seq cells, along with population annotations, are shown to the right.

**Figure S3:**
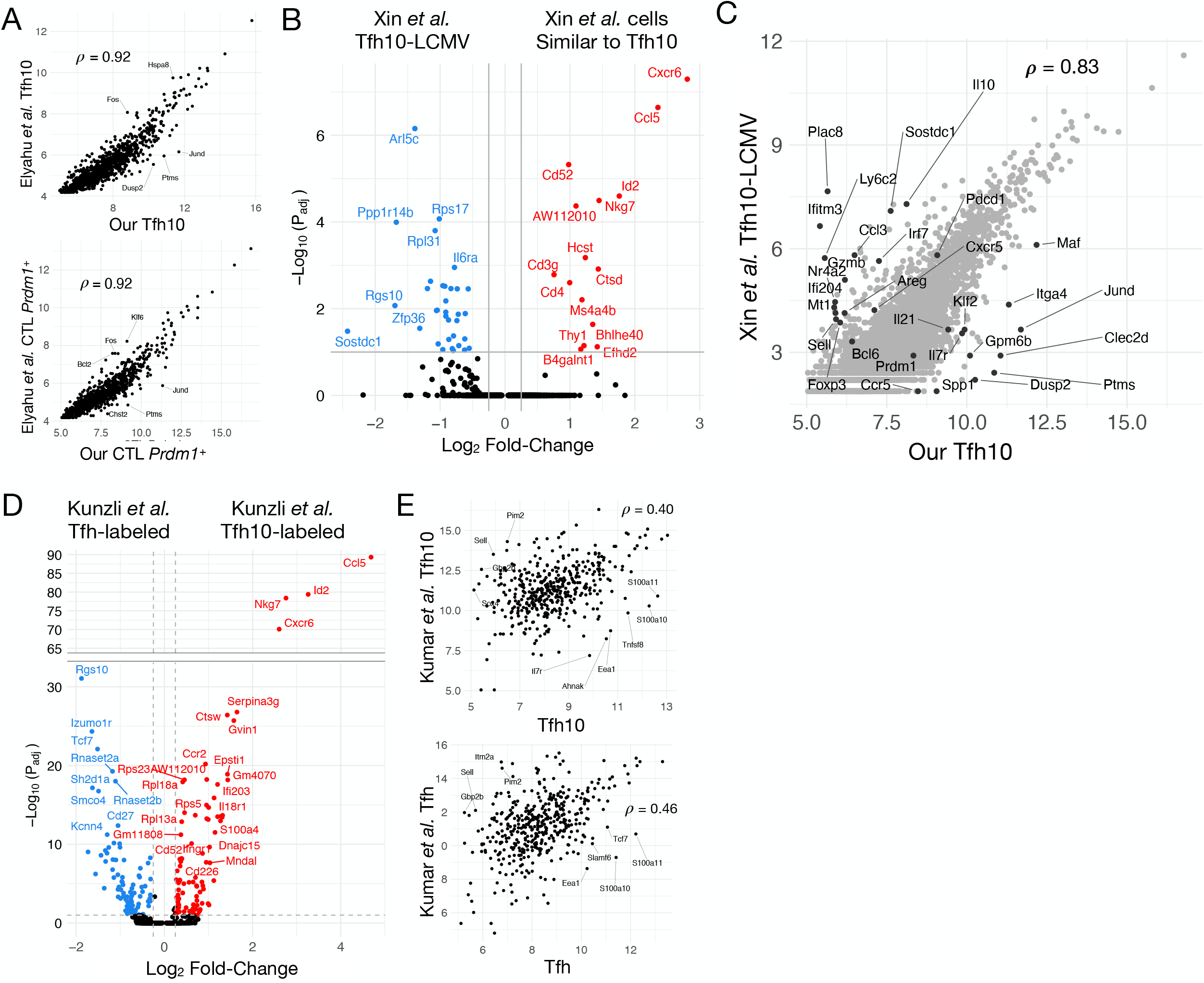
Cross-comparative analyses of other CD4^+^TM scRNA-seq studies, related to Figure 2. **(A)** Comparison of pseudobulk gene expression (DESeq2 VST-normalized counts) between our Tfh10 (top) and *Prdm1*^+^CD4^+^CTL (bottom) populations and those identified by label transfer to Elyahu *et al*. Pearson correlation is shown. **(B)** Differentially expressed genes (|Log2FC| > 0.25, FDR=10%) between cells originally annotated as Tfh10-LCMV and cells mapping to our Tfh10 population based on label transfer from (Xin et al. 2018). **(C)** Comparison of gene expression (DESeq2 VST-normalized counts) of our Tfh10 versus cells originally annotated as Tfh10-LCMV from (Xin et al. 2018). Pearson correlation is shown. **(D)** Differentially expressed genes (|Log_2_FC| > 0.25, FDR=10%) between cells originally annotated as Tfh versus cells that mapped to our Tfh10 based on label transfer to (Künzli et al. 2020). **(E)** Comparison of pseudobulk gene expression between our Tfh10 (top) and Tfh (bottom) populations and those identified by label transfer to human cells (Kumar et al. 2021).

**Figure S4:**
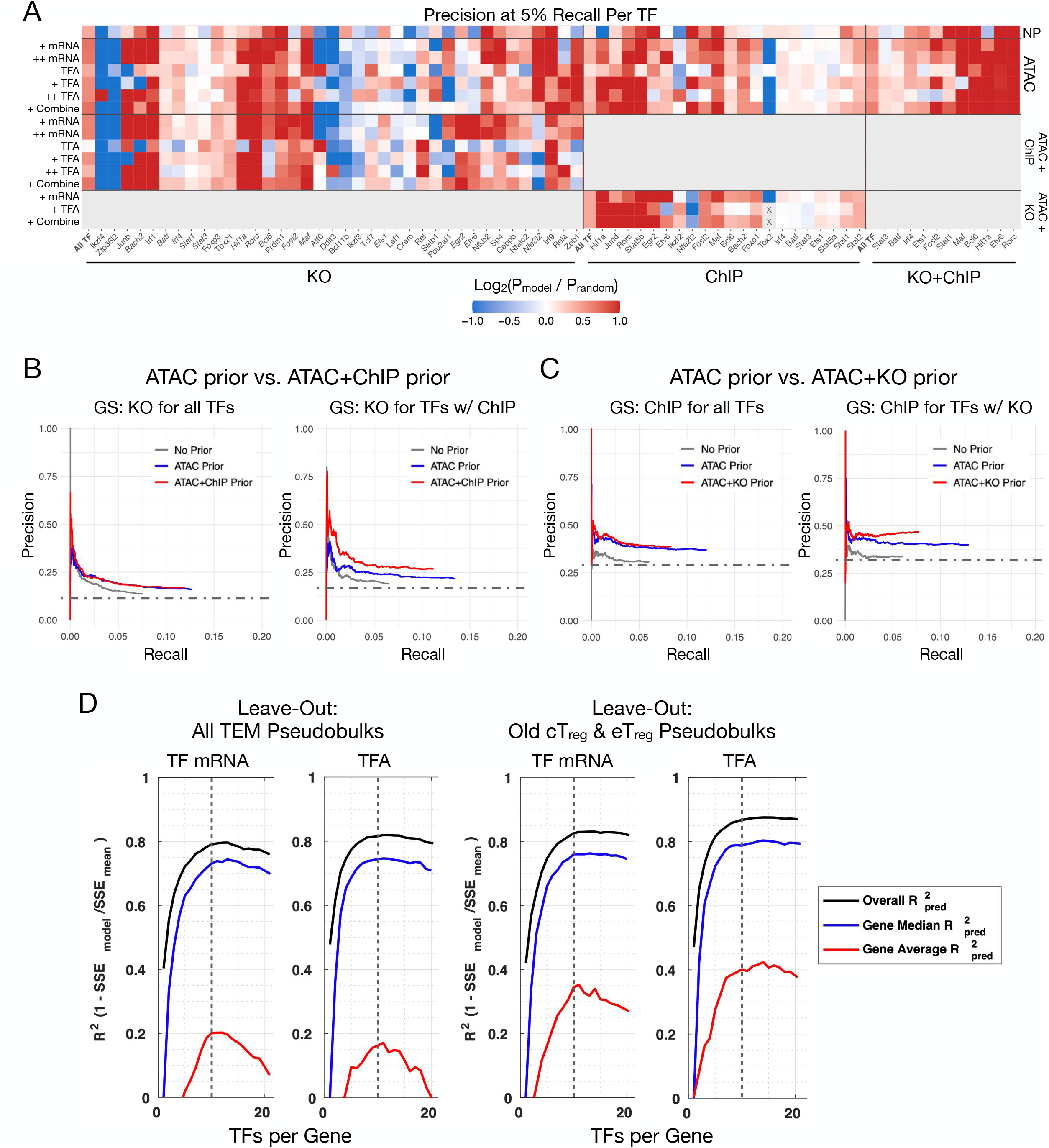
Benchmarking of the CD4^+^TM GRN, related to Figure 4. **(A)** TF-specific GRN performance. Precision at 5% recall, relative to random, of gold standard (**GS**) TF-gene interactions for GRNs inferred with different model parameters. (Precision is the fraction of GRN model predictions supported by the GS (KO and/or ChIP interactions), while recall is the fraction of the GS interactions predicted by the GRN.) We tested inference parameters for TFA estimation method (TF mRNA versus prior-based TFA, **Equation 3**), level of prior reinforcement (+=moderate, bias=0.5; ++=high, bias=0.25, **Equation 2**), and source of prior information (NP, no prior; ATAC, ATAC-based prior; ATAC+ChIP, ATAC-based prior refined by ChIP-seq-derived interactions, see **Methods**). The “combined” GRNs integrate TF mRNA and prior-based TFA networks (see **Methods**). Along the bottom are TFs in the TF knockout (**KO**), ChIP, and KO+ChIP gold standards **(GS).** Grayed-out regions correspond to improper comparisons (e.g., ChIP data was used in the prior and therefore could not be evaluated using the ChIP data in the gold standard as well). Squares are marked with “X” if the model did not reach 5% recall, for the given TF. **(B,C)** Supplementing the ATAC prior with **(B)** TF ChIP-seq or **(C)** TF KO leads to a performance improvement relative to an ATAC-only or no prior network. **(D)** GRN quality was also assessed by out-of-sample gene expression prediction (see **Methods**). We built GRNs excluding either (left) all TEM populations or (right) old cT_reg_ and eT_reg_ populations, and then evaluated the GRN based on gene expression prediction for the left-out (“test”) cell type-conditions. Predictive performance (see **Methods**) was quantified with overall 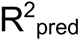 or median or averaged gene 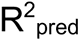 as a function of model size (TFs/gene). Because predictive performance falls off or plateaus at ∼10 TFs/gene, we selected 10 TFs/gene as the model confidence cutoff for our final CD4^+^TM GRN.

**Figure S5:**
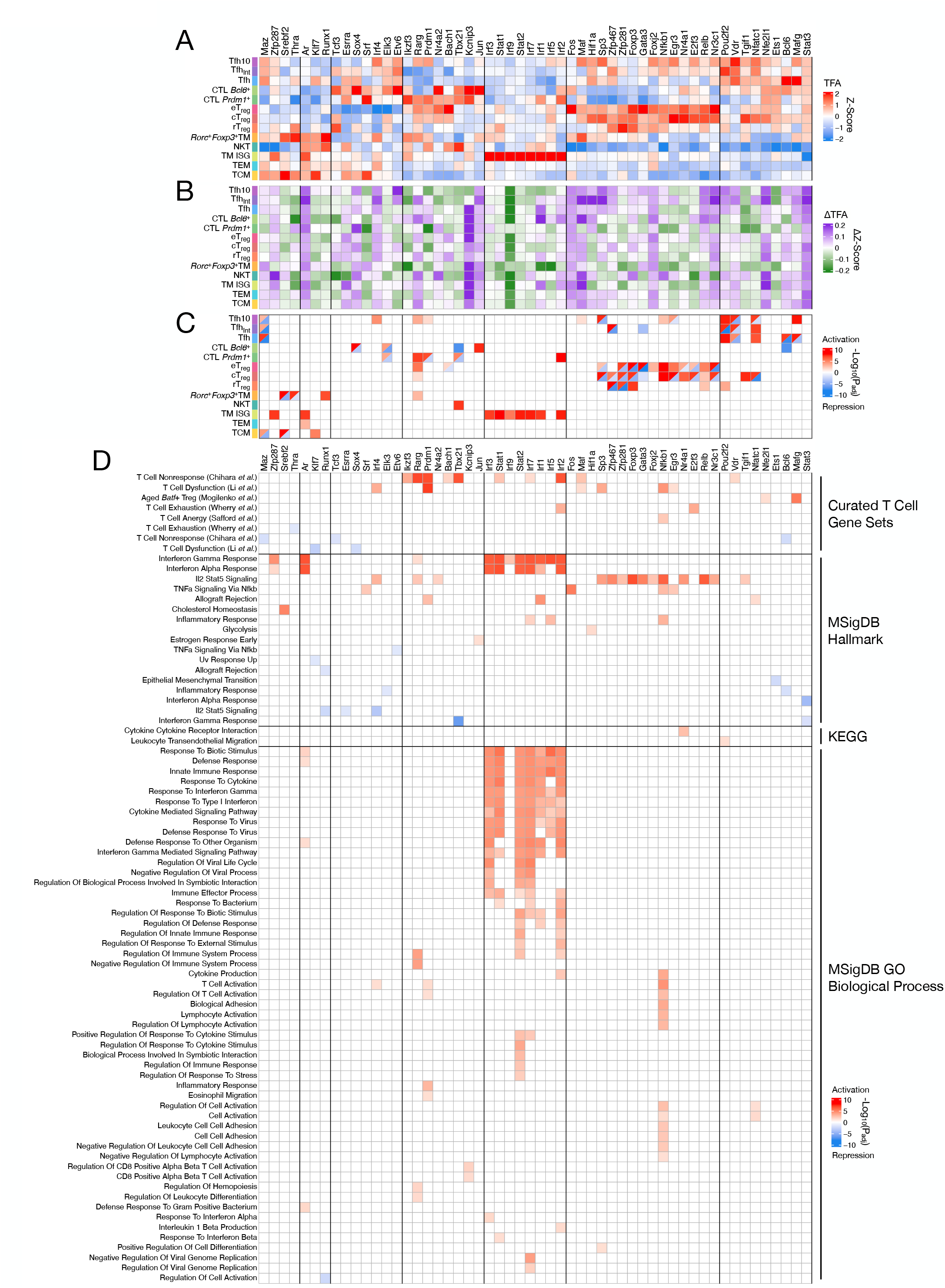
Functional predictions for TFs based on GSEA of their target genes, related to Figure 4. **(A)** Estimated protein TF activities, averaged across age groups and biological replicates, based on the final GRN (see **Methods**, **Equation 3**). **(B)** Differences in TF activities between young and old populations within each T-cell subset. Green indicates higher activity in young, and purple indicates higher activity in old cells. **(C)** “Core” TF annotations; red indicates TF activator, while blue indicates repressor activity. **(D)** GSEA of inferred TF gene targets from our GRN of CD4^+^TM cells. Enrichment of gene pathways in gene targets activated (red) or repressed (blue) by corresponding TF (hypergeometric CDF, FDR=5%).

**Figure S6:**
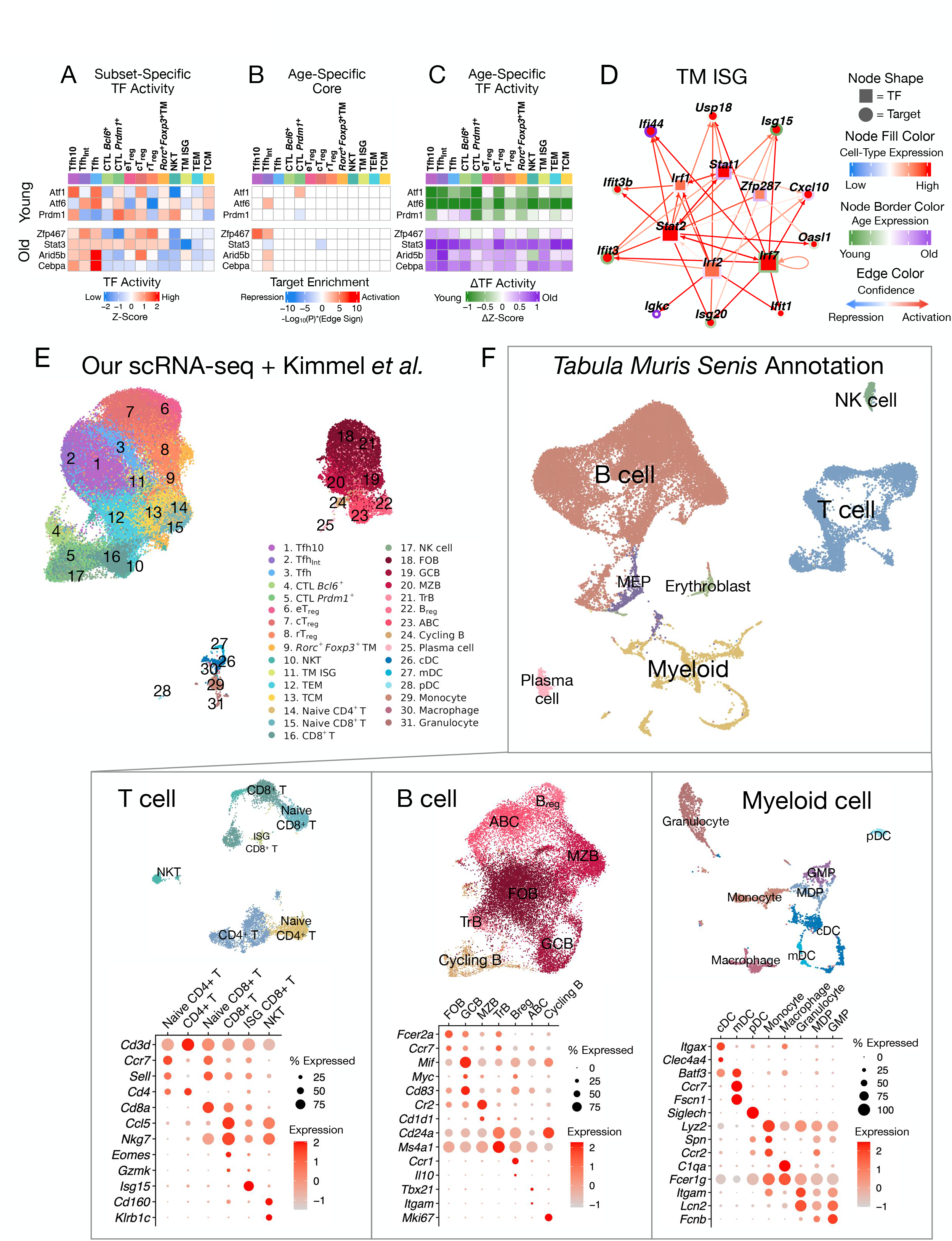
Subset-specific regulators of age-dependent gene expression, the ISG TM core GRN and pan-cell atlas annotations, related to Figures 5-7. **(A-C)** “Core” TFs controlling age-dependent gene signatures in young (top) and old (bottom) populations. **(A)** Estimated protein TF activities, averaged across age groups and biological replicates, based on the final GRN (see Methods, Equation 3). **(B)** “Core” age-specific colored according to whether they promote age-dependent signature within a T-cell population via activation (red) or repression (blue). (C) Differences in TF activities between young and old populations within each T cell subset. Green indicates higher activity in young, and purple indicates higher activity in old cells. (D) Selected GRN interactions between core TFs and signature genes for the TM ISG population (see Fig. 6 for core TF networks of other CD4^+^TM populations). Node shape distinguishes between TFs (square) and targets (circle). Node fill color indicates subset-specific gene expression, and node border color indicates age-dependent gene expression within that population. Edge color indicates either activation (red) or repression (blue) of gene expression. (E) UMAP visualization of our scRNA-seq dataset integrated with age-matched, splenic cells from (Kimmel et al. 2019). (F) Reannotation of young and old splenic cells from the *Tabula Muris Senis* aging pan-cell atlas, including refined annotation of T cell, B cell, and myeloid subpopulations (see Methods). Dot plots show expression of select markers used to annotate subclusters. Abbreviations: FOB, follicular B cell; GCB, germinal center B cell; MZB, marginal zone B cell; TrB, transitioning B cell; Breg, regulatory B cell; ABC, age-associated B cell; cDC, conventional dendritic cell (DC); mDC, mature DC; pDC, plasmacytoid DC.

**Figure S7:**
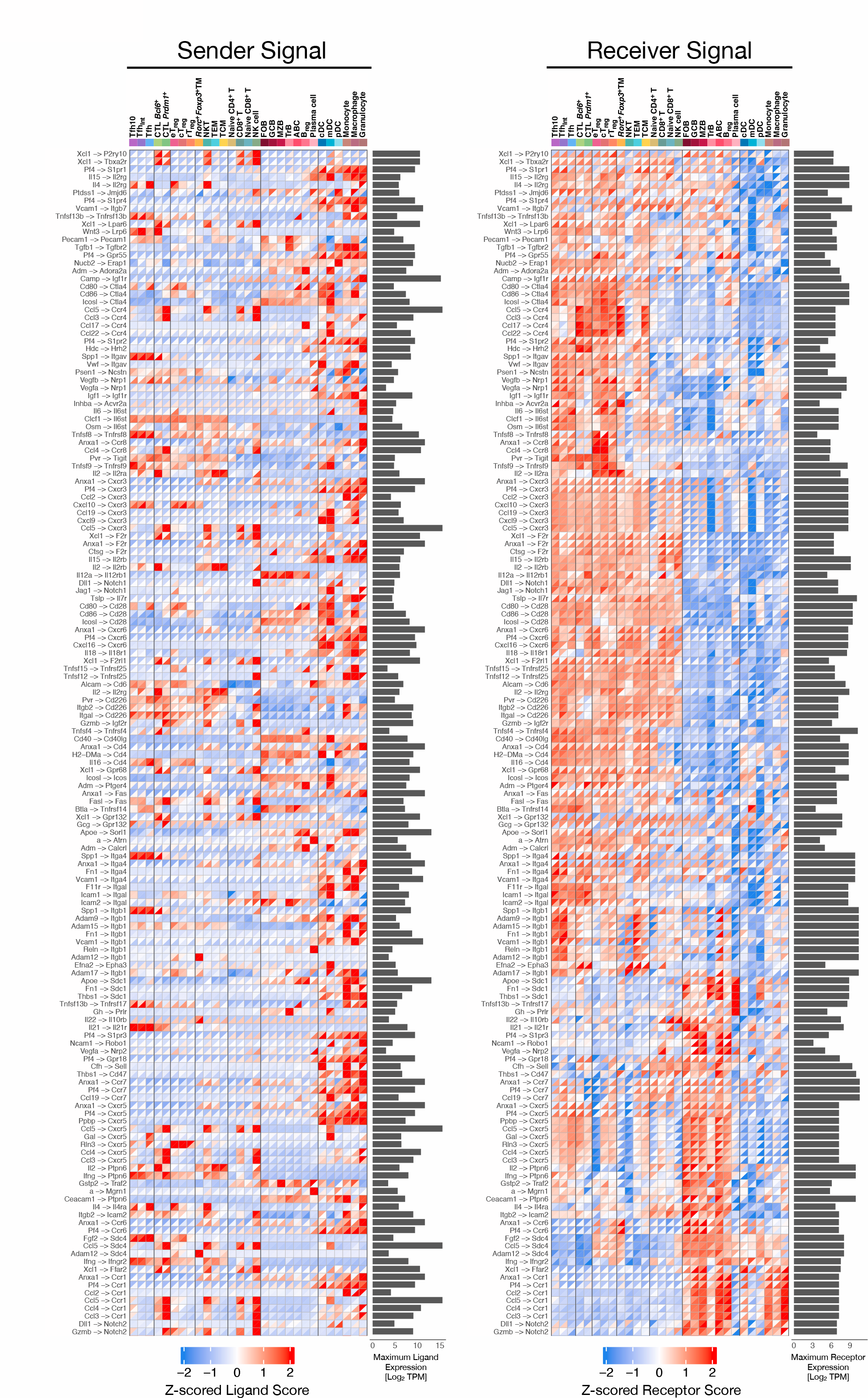
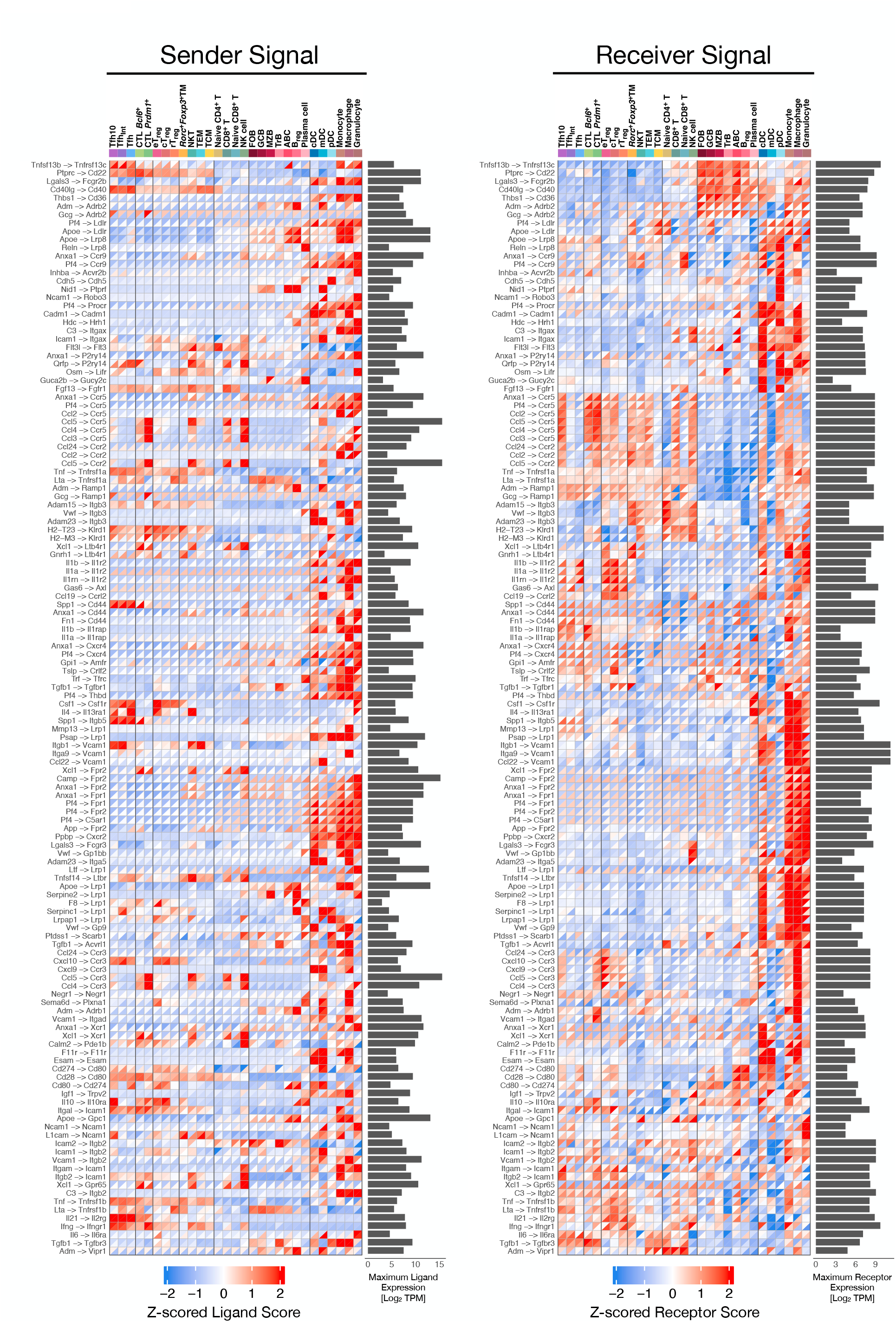
Pan-immune intercellular signaling prediction for young and aged spleen, related to Figure 7. Cell-signaling interactions inferred from our scRNA-seq CD4^+^TM dataset integrated with age-matched splenic cells from *Tabula Muris Senis* (Almanzar et al. 2020). Cell-type-specific sender and receiver signals are shown for young (top triangle) and old (bottom triangle) cells within each cell type. Sender signal corresponds to NicheNet scaled ligand scores, while receptor signals correspond to scaled receptor scores. These predictions are Z-scored across all cell types and age groups. To complement relative ligand and receptors scores, the maximum ligand and receptor expression (gray bar graph; Log2 transcripts per million (**TPM**)) enables comparison of absolute transcript abundance per ligand or receptor. Abbreviations: FOB, follicular B cell; GCB, germinal center B cell; MZB, marginal zone B cell; TrB, transitioning B cell; B_reg_, regulatory B cell; ABC, age-associated B cell; cDC, conventional dendritic cell (DC); mDC, mature DC; pDC, plasmacytoid DC.

## Supplemental Table Legends

**Table S1: Gene Set Enrichment Analysis (GSEA) of population- and age-dependent gene pathway signatures, related to** **Figure 3**. Cell-type-specific and age-dependent gene pathway enrichments (FDR=5%) are shown for curated T-cell-related gene sets as well as gene sets from databases GO, KEGG, and MSigDB.

**Table S2: T cell population-specific signature genes.** For each T-cell population, we identified a set of “signature genes” exhibiting high or low expression relative to other populations. For a given population, we defined an upregulated signature as genes more highly expressed (Log2(FC) > 1, FDR=10%) in that subset relative to at least one other population and not decreased (Log2(FC) < -1, FDR=10%) relative to any other subset. Similarly, for each population, we defined a downregulated signature as genes less expressed in that population relative to at least one other population and not increased relative to any other population.

**Table S3: Age-dependent signature genes.** For each T-cell population, we identified age-dependent “signature genes”, differentially expressed (**DE**) between young and old cells (|Log2(FC)| > 0.25, FDR=10%). We additionally required that each DE gene be expressed in at least 5% of cells in the corresponding subset. Genes with a positive fold-change have higher pseudobulk expression in old cells, and genes with a negative fold-change have higher pseudobulk expression in young cells.

**Table S4: Gold standard (GS) gene regulatory network.** We curated networks of TF-gene interactions in CD4^+^ T cells derived from TF ChIP-seq (“TF Binding GS”) and/or TF knockout followed by RNA-seq (“TF Perturbation GS”). We list metadata for the published samples we processed (see **Methods**) and included in the GS networks. The “Combined GS” network combines interactions for TFs, supported by both binding and perturbation experiments. The interaction signs are based on the TF perturbation data.

**Table S5: Final gene regulatory network (GRN).** Using a prior network informed by scATAC-seq, TF ChIP-seq and perturbation data, the final GRN network combines TF-target gene predictions from networks based on TF mRNA and prior-based TFA by taking the maximum interaction confidence between networks. We restricted the size of the final GRN to an average of 10 TFs per gene based on out-of-sample gene expression prediction (see **Methods**).

**Table S6: Pseudobulk gene expression and chromatin accessibility.** Raw and DESeq2 variance stabilizing transformation-normalized gene expression and chromatin accessibility derived from pseudobulk T-cell populations. Normalized values are batch-corrected with ComBat. The naming convention of the columns is <CELL-type>_<AGE>_<BIOLOGICAL Replicate>.

**Table S7: Prior gene regulatory networks (GRNs).** ATAC-based and final prior GRNs used during benchmarking and final GRN construction, respectively. “ATAC prior” consists of TF-gene interactions supported by putative TF binding sites (**TFBS**) in regions of accessible chromatin proximal to target genes. “Final prior” consists of the ATAC prior with interactions replaced by those supported by gold standard ChIP-seq and/or TF knockout RNA-seq experiments. Interaction weights are Frobenius-normalized per TF.

